# Deletion of *Vhl* in *Dmp1*-expressing cells causes microenvironmental impairment of B cell lymphopoiesis

**DOI:** 10.1101/2021.09.10.459794

**Authors:** Betsabel Chicana, Nastaran Abbasizadeh, Christian Burns, Hanna Taglinao, Joel A. Spencer, Jennifer O. Manilay

**Affiliations:** Department of Molecular and Cell Biology, School of Natural Sciences, University of California, Merced, Merced, CA, USA; Quantitative and Systems Biology Graduate Program, University of California, Merced, Merced, CA, USA; Department of Bioengineering, School of Engineering, University of California, Merced, Merced, CA, USA; Bioengineering Graduate Program, University of California, Merced, Merced, CA, USA

**Author notes:** **Joint Senior Authors (J.A.S and J.O.M)**. **Correspondence to (J.A.S.) and (J.O.M.)**.

## Abstract

The contributions of skeletal cells to the processes of B cell development in the bone marrow (BM) have not been completely described. The von-Hippel Lindau protein (VHL) plays a key role in cellular responses to hypoxia. Previous work showed that *Dmp1*-Cre;*Vhl* conditional knockout mice (*Vhl*cKO), which delete *Vhl* in late osteoblasts and osteocytes, display dysregulated bone growth and reduction in B cells. Here, we investigated the mechanisms underlying the B cell defects using flow cytometry and high-resolution imaging. In the *Vhl*cKO BM, B cell progenitors were increased in frequency and number, whereas Hardy Fractions B-F were decreased. *Vhl*cKO Fractions B-C cells showed increased apoptosis and quiescence. Reciprocal BM chimeras confirmed a B cell-extrinsic source of the *Vhl*cKO B cell defects. In support of this, *Vhl*cKO BM serum contained reduced CXCL12 and elevated EPO levels. Staining of *Vhl*cKO B cells with an intracellular hypoxic marker indicated the natural existence of distinct B cell microenvironments that differ in local oxygen tensions. Additionally, intravital and ex vivo imaging revealed *Vhl*cKO BM blood vessels with increased diameter, frequency, volume, and a diminished blood-BM barrier. Our studies identify novel mechanisms linking altered bone homeostasis with drastic BM microenvironmental changes that dysregulate B cell development.

## INTRODUCTION

The mechanisms by which changes in bone homeostasis affect immune development in the bone marrow (BM) are not fully understood (1–4). A detailed understanding of how bone microenvironments affect immune cell development and function could provide strategies towards novel therapeutic approaches to immune deficiencies. B cells produce antibodies (Abs), which are crucial for a robust adaptive immune response. B cells are generated from hematopoietic stem cells (HSCs) in the liver during fetal life, and in the BM in the adult (5). B cell development in the BM occurs in a series of defined stages that rely on growth factors that are produced by osteolineage derived cells in the BM microenvironment, such as mesenchymal stem cells (MSCs) and osteoblasts (OBs) (1).

The von-Hippel Lindau protein (VHL) regulates hypoxia-inducible factor (HIF) degradation, which is involved in cellular adaptation to low oxygen environments (6). When HIF1α accumulates in normoxic conditions, it travels to the nucleus to activate over 100 hypoxia-inducible target genes (7). VHL is expressed ubiquitously in many cell types, and global deletion of the *Vhl* gene results in embryonic lethality, so conditional knockout approaches are necessary to investigate the cell-specific roles of VHL in specific microenvironments. Conditional-deletion of *Vhl* in OBs and in hematopoietic progenitors have demonstrated a role for VHL in these cell types (8, 9). OBs support B cell development, and also mature into osteocytes (OCYs). The role of HIF and its regulation on the immune system has been extensively reviewed, but the mechanism of cell-intrinsic and cell-extrinsic VHL on specific immune cell lineages has not fully been addressed (10).

The BM microenvironment manifests hypoxic heterogeneities in a spatio-temporal manner (11), however the implications of these oxygen tension (pO_2_) differences on hematopoiesis are not well characterized. Hypoxia slows the processes of angiogenesis and osteogenesis during fracture healing and bone formation, but also promotes OB differentiation into OCYs (12), and can stimulate osteoclast formation (13). Studies have shown HIF stabilization as a therapeutic option for treating bone fractures (14, 15) and osteoporosis (16–18), but the underlying molecular mechanism remains poorly understood. VHL plays an important role regulating HIF expression, and disruption of VHL in bone cells leads to improper bone homeostasis. VHL depletion in osteochondral progenitor cells and osteocalcin-positive OBs leads to an increase in bone mass through an increase in OB number (7, 19). Furthermore, disrupting VHL in OBs induces expression of β-catenin, revealing the mechanism by which VHL/HIF pathway promotes bone formation through the Wnt pathway (7, 20, 21). Altogether, these studies of *Vhl* deletion in osteolineage cells have not examined the cell-extrinsic effects of these changes on the immune cells residing in the BM.

The BM contains specialized microenvironments that maintain blood cells and supply factors required for their development and maintenance. Perivascular stromal cells, osteoprogenitor cells, endothelial cells (ECs), MSCs, OBs and OCYs are critical B cell “niches” and are all cells that support B cell development. Collectively, these cells provide essential cytokines for B cell development which include CXC-chemokine ligand 12 (CXCL12), FLT3 ligand (FLT3L), IL-7, stem-cell factor (SCF) and receptor activator of nuclear factor-κB ligand (RANKL) (1, 22–31). Development of B cells starts at the pre-pro-B cell (Fraction A) stage where they are near OBs and CXCL12+ reticular cells, and as B cells continue to mature to the pro-B cell stage (Fractions B-C), they move closer to IL-7 expressing cells; the CXCL12/IL7 levels in the niche is crucial to sustain B cell development (4, 27–29, 31, 32). Furthermore, the BM contains a dense vascular network and vascular sinuses creating the perivascular region, which provides a niche where B cells are known to develop and reside (33). During aging, vascular density decreases in many tissues due to impaired angiogenesis caused by EC dysfunction. Vascular “hyperpermeability” also increases with age, via changes in ECs lining the blood vessel wall, disrupting the blood-BM barrier (34–36). The role of the vasculature and regulation of vessel permeability in hematopoiesis, especially in B cell development, remains unknown.

To understand how changes in bone homeostasis may affect immune cell development, our laboratories utilized *Dmp1*-Cre;*Vhl* conditional knockout mice (*Vhl*cKO), in which *Vhl* is deleted primarily in late OBs and OCYs. In the *Vhl*c*KO* bones, the number of hematopoietic cells is severely reduced, and B cell development is stunted (37). Here, we provide evidence for molecular, cellular and structural changes in the *Vhl*cKO BM niche that adversely affect B cell development in a cell-extrinsic manner, such as reduction in key niche cells, decreased production of B cell supporting cytokines, and structural changes in the BM vasculature. These studies reveal novel molecular mechanisms by which *Vhl* deletion in *Dmp1*-expressing cells affect B cell niches.

## MATERIALS AND METHODS

### Study design

A G*Power statistical (38) power analysis (α=0.05 and power of 0.95) based on B cell developmental data and BM cellularity determined that a minimum of n=7 mice per group was needed for our studies. The total sample size for each experiment was >7 performed in three independent experiments. Age-matched mice of both sexes *Vhl*cKO and control mice (C57BL/6 wild type and *Vhl*-floxed (*Vhl^fl/fl^*, *Dmp1-Cre-*negative mice) were used and no sex-specific differences in B cell development or other relevant characteristics to our studies were detected. Student’s t-test and nonparametric Bonferroni-corrected Mann-Whitney U-test was used to test differences between mean and median values with Graph-Pad Prism and were considered significant if p<0.05.

### Experimental Animals

Mice on the C57Bl/6 background were used. B6N.FVB-Tg1Jqfe/BwdJ (*Dmp1*-Cre) (39) and B6.129S4(C)-Vhl tm1Jae/ J (*Vhl^fl/fl^*) (40) were purchased from The Jackson Laboratory. These two lines of mice were crossed to generate *Vhl* conditional knockouts in *Dmp1*-expressing cells (*Vhl*cKO). Genotyping was confirmed following protocols from Jackson Laboratory for JAX Stock #021047 and #012933. Mice were housed under specific pathogen-free conditions in the University of California, Merced’s vivarium with autoclaved feed and water, and sterile microisolator cages. The University of California Merced Institutional Animal Care and Use Committee approved all animal work.

### Bone marrow transplantation

Recipient mice were 10 weeks of age at the time of transplantation. Whole bone marrow WT (CD45.1+) donor cells (1×10^6^) were injected retro-orbitally into lethally irradiated (1000 rads using a Cesium-137 source, JL Shepherd and Associates, San Fernando, CA, USA) recipient *Vhl*cKO (CD45.2+) mice under isoflurane anesthesia. Reciprocal *Vhl*cKO→WT chimeras were also prepared. Animals were supplemented with neomycin in the drinking water for 14 days post-transplant as described (41).

### Sample Collection: Bone Marrow, Peripheral Blood, Spleen and Serum

#### Bone marrow collection

Mice were euthanized by the inhalation of carbon dioxide followed by cervical dislocation. Femurs and tibias were dissected, and muscles were removed. To release the BM, bones were crushed with a mortar and pestle in M199+ (M199 with 2% FBS). BM cells were collected into 15mL conical tubes after being rinsed away from bone chips with M199+, resuspended by trituration, filtered through 70-micron nylon mesh into a 50 mL conical tube, and centrifuged for 5 mins at 1500 rpm and at 4°C. Cell pellets were resuspended and treated with ACK lysis buffer to remove erythrocytes. Cells treated with ACK were washed and resuspended in M199+. Cell counts were obtained using a hemocytometer and Trypan Blue staining to exclude dead cells.

To collect BM serum, femurs were cleaned of any muscle tissue and the epiphyses were cut off and discarded. The bone shaft was then placed into a 0.2 mL tube in which a hole was introduced using a needle. Thirty μL of 1x phosphate buffered saline (PBS) was placed on the top end of the bone shaft, using a 25g needle, and then the tube containing the bone was placed into a 1.5 ml microcentrifuge tube and centrifuged for 30 seconds at 15,000rpm. Serum supernatant was collected and stored at −80C until analysis.

#### Peripheral blood collection

Mice were heated under a heat lamp to increase blood circulation and then restrained. Blood collection was performed via tail bleeds by making an incision with a scalpel blade over the ventral tail vein. No more than ten drops were collected (<0.5 mL) in a 1.5 ml Eppendorf tube with 50 uL of heparin. To obtain blood serum, blood was collected in 1.5 ml tubes without heparin and allowed to clot for 30 minutes at room temperature. The samples were then centrifuged for 10 minutes at 4000 rpm at 4°C. Blood serum was collected and stored at −80°C until the day of analysis.

#### Spleen cell collection

Dissected spleens were processed and mashed in 1 mL of ACK lysis buffer in a petri dish for no more than one minute. Five mL of M199+ were added into the dish to dilute the ACK lysis buffer and to stop red cell lysis. Spleen cells were aspirated into a 5mL syringe to create single cell suspensions by passing the cells through the syringe several times then filtering through a 70-micron nylon mesh into a 15 mL conical tube. Cells were centrifuged at 2000 rpm at 4°C for 3 minutes. Cell pellets were finger vortexed and resuspended with 5 mL of M199+. Live cell counts were determined using a hemocytometer and Trypan Blue staining.

### Quantification of cytokines

Cytokine measurements were performed using a customized bead-based multiplex (13-LEGENDplex assay) from Biolegend, Inc. with the analytes IL-3, IL-5, IL-6, IL-7, IL-15, IL-34, M-CSF, TPO, GM-CSF, LIF, EPO, CXCL12, SCF for the analysis of BM serum and peripheral blood serum of *Vhl*cKO and control mice. Concentrations of cytokines were determined from samples following manufacturer’s instructions and software.

### Flow cytometry analysis and antibodies

Cells were stained for flow cytometry and included a pre-incubation step with unconjugated anti-CD16/32 (clone 93) to block Fc receptors as previously described (41, 42). Samples were stained with antibodies listed in **Supplemental Table 1**. For viability staining, DAPI or PI was used. Single color stains were used for setting compensations and gates were determined with fluorescent-minus one controls, isotype-matched antibody controls, or historical controls. Intracellular staining of Ki67 was performed using the eBioscience™ Foxp3 / Transcription Factor Staining Buffer Set following the manufacturer’s instructions. Apoptosis staining was performed using Biolegend Annexin V Apoptosis Detection Kit with 7AAD. Flow cytometry data was acquired on the BD LSR II. The data was analyzed using FlowJo Software version 10.7.1.

### Preparation of long bones for imaging

To label blood vessels, mice were injected with fluorescent antibodies (**Supplemental Table 1**) through the retro-orbital venous sinus. After 20 minutes of incubation, intracardial perfusion was performed with 1X PBS following by cold and fresh 4% paraformaldehyde (PFA). Subsequently, femurs were harvested and fixed in the 4% PFA for 30 minutes, at 4° C. The bones were then washed with 1X PBS, immersed in 30% sucrose for 1 hour, frozen in optimal cutting temperature (OCT) compound and kept at – 80° C. Samples were shaved using a cryostat (LEICA CM1860) equipped with a high-profile blade (Leica; 3802121).

To optically clear long bones, a modified uDISCO clearing protocol was used (43). After intracardial perfusion as described above, long bones were immersed in 4% PFA overnight and put through a series of *tert*-butanol dehydration steps at 30% (4 hours), 50% (4 hours), 70% (overnight), 80% (4 hours), 90% (4 hours), and 100% (overnight). Next, long bones were incubated in dichloromethane (DCM) for 40 minutes and then placed in Benzyl Alcohol Benzyl Benzoate - DL-alpha-tocopherol (BABB-D4) for 3-4 hours. BABB-D4 is prepared by mixing Benzyl Alcohol + Benzyl Benzoate at the ratio of 1:2, adding diphenyl ether (DPE) to the BABB solution (1:4) and ultimately DL-alpha-tocopherol (Vitamin E) with the ratio of 1:25 to decrease fluorescence quenching. Cleared femurs were mounted in a custom glass chamber filled with BABB-D4 and sealed with solvent-resistant silicone gel (DOWSIL™ 730) (43).

### Two-photon microscopy

Imaging was performed with a custom-built two-photon video-rate microscope (Bliq Photonics) equipped with two femtosecond lasers (Spectra Physics; Insight X3, Spectra Physics; MaiTai eHP DS). During intravital imaging, the Spectra Physics Insight X3 and Maitai laser wavelengths were tuned to 840 nm and 1040 nm, respectively, and for ex vivo imaging only the Insight X3 was tuned to 1220 nm. Three fluorescent channels were acquired (520-535 nm, 590-636 nm, and 679-741 nm). For all two-photon imaging, a 25x water immersion objective (Olympus; XLPLN25XWMP2) with 1.05 numerical aperture was used to image a 317 μm by 159 μm field of view. Videos were recorded at 30 frames per second and images were generated by averaging of 30 frames from the live video mode.

For in vivo imaging of calvarial bone marrow, mice were anesthetized with isofluorane (3-4% induction, 1.5% maintenance at 1L/min) and the top of the head shaved. The skin was cleaned with 70% alcohol wipes before surgery. The mouse was placed on a heating pad and secured in a custom head mount. An incision was made along the sagittal and lambda suture of the skull and the skin retracted to expose the calvarial bone as previously described (11, 44). The secured mouse was then placed on the microscope stage for two-photon microscopy(11, 44). In order to measure BM blood vessel permeability, leakage and flow velocity in the calvaria BM during in vivo imaging, 70 kDa Rhodamine-B-Dextran (ThermoFisher; D1841) was injected retro-orbitally while the mouse was on the stage.

For ex vivo imaging, optically cleared long bones were mounted in a chamber sealed with solvent-resistant silicone gel and shaved long bones were mounted on a wet sponge to prevent the sample from drying during imaging. Slides were imaged with similar acquisitions settings as the in vivo imaging.

### Image quantification

For in vivo image analysis, image processing and permeability/leakage measurements were performed with Fiji (ImageJ 1.53k) and BM blood flow velocity was quantified with custom scripts in MATLAB (2020a). To measure permeability in the calvaria, live two-photon microscopy video was recorded for the first 30 seconds after Rhodamine B Dextran was injected. The blood vessel permeability was calculated based on the change in fluorescence intensity outside of blood vessels over time as previously described (45, 46). For leakage measurements, z-stacks (2 μm step size) were recorded randomly around the calvarium BM 10 minutes after injection. Leakage values were calculated by dividing the fluorescence intensity of the perivascular space adjacent to a vessel by the fluorescence intensity inside the blood vessel. Representative examples of BM leakage were generated by taking maximum intensity projections (MIPs) of BM regions with image contrast/enhancement applied. Blood flow velocity was calculated by recording 30 second videos of blood flow in the BM calvaria and then utilizing the Line Scanning Particle Image Velocimetry (LSPIV) method implemented in a custom MATLAB script to calculate blood flow velocity as previously described (47, 48). ImageJ (ImageJ 1.53k) was used to adjust video and image contrast for figure presentation.

To generate a depth-dependent profile of vessel diameter in long bones, measurements were taken at 0-30 μm (shallow BM), 75-105 μm (middle BM), and 150-180 μm (deep BM) below the endosteum. To measure vascular density, image brightness/contrast was first adjusted in Fiji (ImageJ 1.53k) and then images were converted to binary. Next, noise reduction was performed via Despeckle, and binary Fill Hole was applied. Finally, using analytical coding developed in Python (3.7.6), the ratio of the total blood vessel pixels to total BM pixels was determined for BM vessel density measurements.

## RESULTS

### *Vhl* deletion in Dmp1-expressing cells dysregulates hematopoiesis

Long bones in *Vhl*c*KO* mice display abnormally high bone mass and density and the BM cavity is severely occluded with bone, accompanied by stunted B cell development (37) (**Figure 1A**). Consistent with this, *Vhl*cKO mice displayed splenomegaly, consistent with extramedullary hematopoiesis (**Supplemental Figure 1A-E).** *Vhl*cKO mice exhibit reduced BM cellularity compared to controls (**Figure 1B**) and analysis of specific hematopoietic cell lineages revealed a decrease in B cells, no change in T cell frequency, and an increase in monocytes and granulocytes in 6-week-old, 10-week-old and 6-month-old mice (**Figure 1C-D**). Similarly, lineage analysis in the spleen at 10 weeks revealed a decrease in B cells, no change in T cells, and an increase in granulocytes that became more prominent as mice aged to 6-months. Monocytes in the *Vhl*cKO spleen at 6-weeks-old were slightly reduced, similar to controls at 10-weeks-old, and were increased at 6-months-old (**Supplemental Figure 1E**). Peripheral blood of the *Vhl*cKO mice showed decreased B cells at all ages examined, whereas monocytes were increased at 10-weeks-old, and granulocytes at 6-months-old only (**Supplemental Figure 1F**). Furthermore, we observed greater proportions of monocytes and granulocytes and an overall reduction in the absolute numbers of all hematopoietic lineages in the BM of *Vhlc*KO mice (**Table 1**).

**Figure 1.**
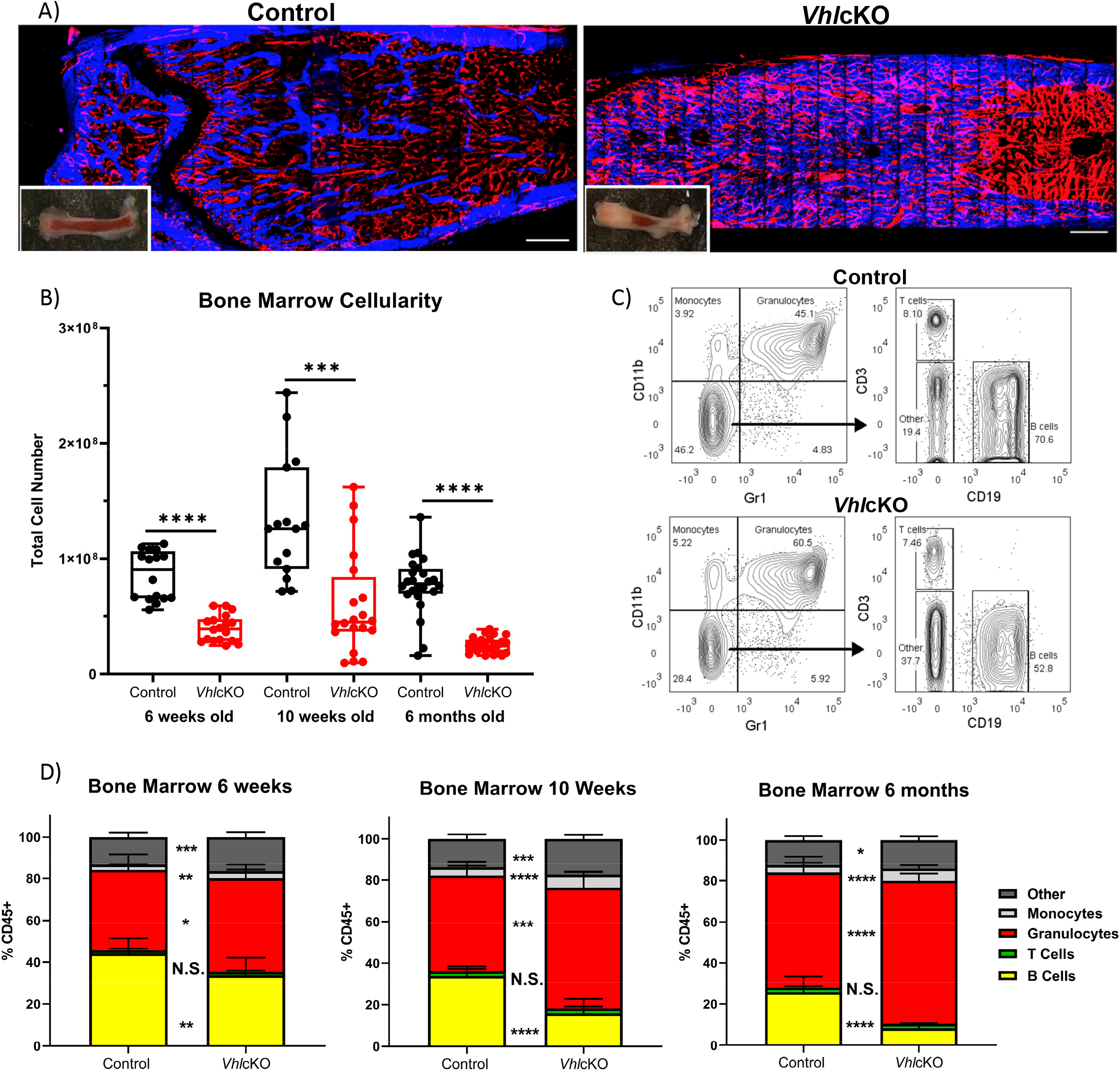
Bone marrow, spleen and peripheral blood lineage cell defects in the *Vhl*cKO mice. A) Macroscopic and ex vivo imaging of the long bones revealed progressive increases in the bone mass in *Vhl*cKO femurs compared with the controls. Inset: photo of the femur. Red: blood vessel (AlexaFluor647 CD31, AlexaFluor647 CD144, AlexaFluor647 Sca-1), Blue: bone (SHG). Scale bar ~100μm; B) bone marrow cellularity, C) representative FACS plots of immune cell lineages; D) frequency analysis of bone marrow lineage cells at 6-weeks of age (left), 10-weeks of age (middle) and 6-month (right). p<0.05*, p<0.01**, p<0.001***, p<0.0001**** two-tailed Student’s t-test.

**Table 1.** Hematopoetic lineage mean ± SD and absolute number. p<0.05*, p<0.01**, p<0.001***, p<0.0001**** two-tailed Student’s t-test.

| | CD45+ population (mean% ± SD) | | | | Absolute Number (mean ± SD) | | | |
| --- | --- | --- | --- | --- | --- | --- | --- | --- |
| Lineage Population | Bone Marrow |  | Spleen |  | Bone Marrow |  | Spleen |  |
| 6 weeks old | Control | *Vhl* cKO | Control | *Vhl* cKO | Control | *Vhl* cKO | Control | *Vhl* cKO |
| B cells | 44.16 ± 7.16 | 33.69 ± 8.49** | 60.71 ± 2.64 | 58.57 ± 3.51 | 2.62E+0.7 ± 7.06E+06 | 8.88E+0.6 ± 4.83E+06**** | 6.35E+0.7 ± 1.28E+07 | 5.71E+0.7 ± 9.18E+06 |
| T cells | 1.45 ± 0.79 | 1.61 ± 0.62 | 23.65 ± 3.56 | 23.49 ± 3.42 | 8.36E+0.5 ± 4.33E+05 | 3.70E+0.5 ± 1.87E+05*** | 2.49E+0.7 ± 6.57E+06 | 2.29E+0.7 ± 4.93E+06 |
| Monocytes | 2.58 ± 0.37 | 3.28 ± 0.74** | 2.55 ± 0.57 | 1.95 ± 0.15** | 1.53E+0.6 ± 3.92E+05 | 7.81E+0.5 ± 3.48E+05**** | 2.66E+0.6 ± 7.87E+05 | 1.90E+0.6 ± 2.96E+05* |
| Granulocytes | 38.46 ± 7.61 | 44.85 ± 6.47* | 2.85 ± 2.55 | 2.28 ± 0.66 | 2.42E+0.7 ± 1.12E+07 | 1.06E+0.7 ± 4.30E+06*** | 2.71E+0.6 ± 2.19E+06 | 2.17E+0.6 ± 5.06E+05 |
| 10 weeks old | Control | *Vhl* cKO | Control | *Vhl* cKO | Control | *Vhl* cKO | Control | *Vhl* cKO |
| B cells | 33.8 ± 4.55 | 15.83 ± 7.01**** | 60.71 ± 7.63 | 46.37 ± 7.53**** | 3.32E+0.7 ± 1.47E+07 | 8.02E+0.6 ± 1.29E+07*** | 1.22E+08 ± 6.02E+07 | 1.18E+08 ± 5.98E+07 |
| T cells | 2.34 ± 1.04 | 2.35 ± 0.95 | 26.67 ± 6.15 | 29.09 ± 4.11 | 2.55E+0.6 ± 1.92E+06 | 1.05E+0.6 ± 1.35E+06* | 6.22E+07 ± 5.28E+07 | 8.10E+07 ± 5.40E+07 |
| Monocytes | 4.09 ± 0.67 | 6.23 ± 1.25**** | 2.50 ± 0.36 | 2.71 ± 0.38 | 4.08E+0.6 ± 2.08E+06 | 2.41E+0.6 ± 1.89E+06* | 5.49E+06 ± 4.12E+06 | 7.02E+06 ± 3.81E+06 |
| Granulocytes | 45.93 ± 6.89 | 58.11 ± 7.78*** | 1.15 ± 0.50 | 6.27 ± 2.93**** | 4.36E+0.7 ± 1.60E+07 | 2.22E+0.7 ± 1.50E+07** | 2.14E+06 ± 9.96E+05 | 1.66E+07 ± 1.46E+07** |
| 6 months old | Control | *Vhl* cKO | Control | *Vhl* cKO | Control | *Vhl* cKO | Control | *Vhl* cKO |
| B cells | 25.74 ± 7.62 | 8.37 ± 1.76**** | 61.38 ± 7.09 | 42.44 ± 3.65**** | 1.48E+0.7 ± 7.35E+06 | 9.73E+0.5 ± 2.56E+05**** | 4.33E+0.7 ± 2.04E+07 | 2.77E+0.7 ± 7.48E+06** |
| T cells | 2.21 ± 0.62 | 2.23 ± 0.38 | 25.55 ± 2.81 | 25.88 ± 2.68 | 1.20E+0.6 ± 5.48E+05 | 2.58E+0.5 ± 5.65E+04**** | 1.66E+0.7 ± 6.08E+06 | 1.70E+0.7 ± 5.08E+06 |
| Monocytes | 3.57 ± 1.31 | 5.98 ± 1.69**** | 1.82 ± 0.47 | 2.60 ± 0.67*** | 1.94E+0.6 ± 9.20E+05 | 6.97E+0.5 ± 2.28E+05**** | 1.19E+0.6 ± 3.95E+05 | 1.74E+0.6 ± 7.95E+05* |
| Granulocytes | 56.01 ± 7.91 | 69.31 ± 3.49**** | 2.83 ± 4.41 | 14.33 ± 3.33**** | 3.08E+0.7 ± 1.26E+07 | 8.22E+0.6 ± 2.11E+06**** | 1.64E+0.6 ± 2.16E+06 | 9.52E+0.6 ± 3.42E+06**** |

### Increased frequencies of hematopoietic progenitor cells in the *Vhl*cKO BM

To further investigate if the defect in hematopoiesis occurred upstream of lineage-committed cells, we analyzed the hematopoietic progenitor compartments in the BM of *Vhl*c*KO* mice. Long-term hematopoietic stem cells (LT-HSCs: LSK, CD150+ CD48-, short term hematopoietic stem cells (ST-HSCs: LSK, CD150-, CD48-), multipotent progenitors (MPP2: LSK, CD150+, CD48+; MPP3: LSK, CD150-, CD48+; and MPP4: LSK, CD150-, Flk2+, CD48+), and common lymphoid progenitors (CLPs: Lineage-, CD127+, cKIT^int^, Sca1^int^) from *Vhl*cKO and control mice were quantified using flow cytometry (**Figure 2A, B**). The results showed an increase in the frequency in LT-HSCs, ST-HSCs, MPP2, MPP3, and CLPs at 6-weeks, 10-weeks and 6-months-old (**Figure 2C**). MPPs are heterogeneous with different lineage-biased potential. MPP2/3 are myeloid-biased while MPP4 are lymphoid-primed (49, 50). In our results, MPP4 frequency was increased starting at 10-weeks-old (**Figure 2C**). These results show that deletion of *Vhl* in Dmp1-expressing cells increases progenitor frequencies and indicates that downstream differentiation of B cells may be blocked. However, examination of MPP4 absolute numbers showed decreased MPP4s in 6-week-old *Vhl*cKO, an increase at 10-weeks-old, and numbers similar to controls at 6-months old. In 10-week-old *Vhl*cKO mice, the absolute numbers of LT-HSCs and MPP3 were increased, whereas at 6-months-old, LT-HSCs and CLPs were decreased (**Figure 2D**).

**Figure 2.**
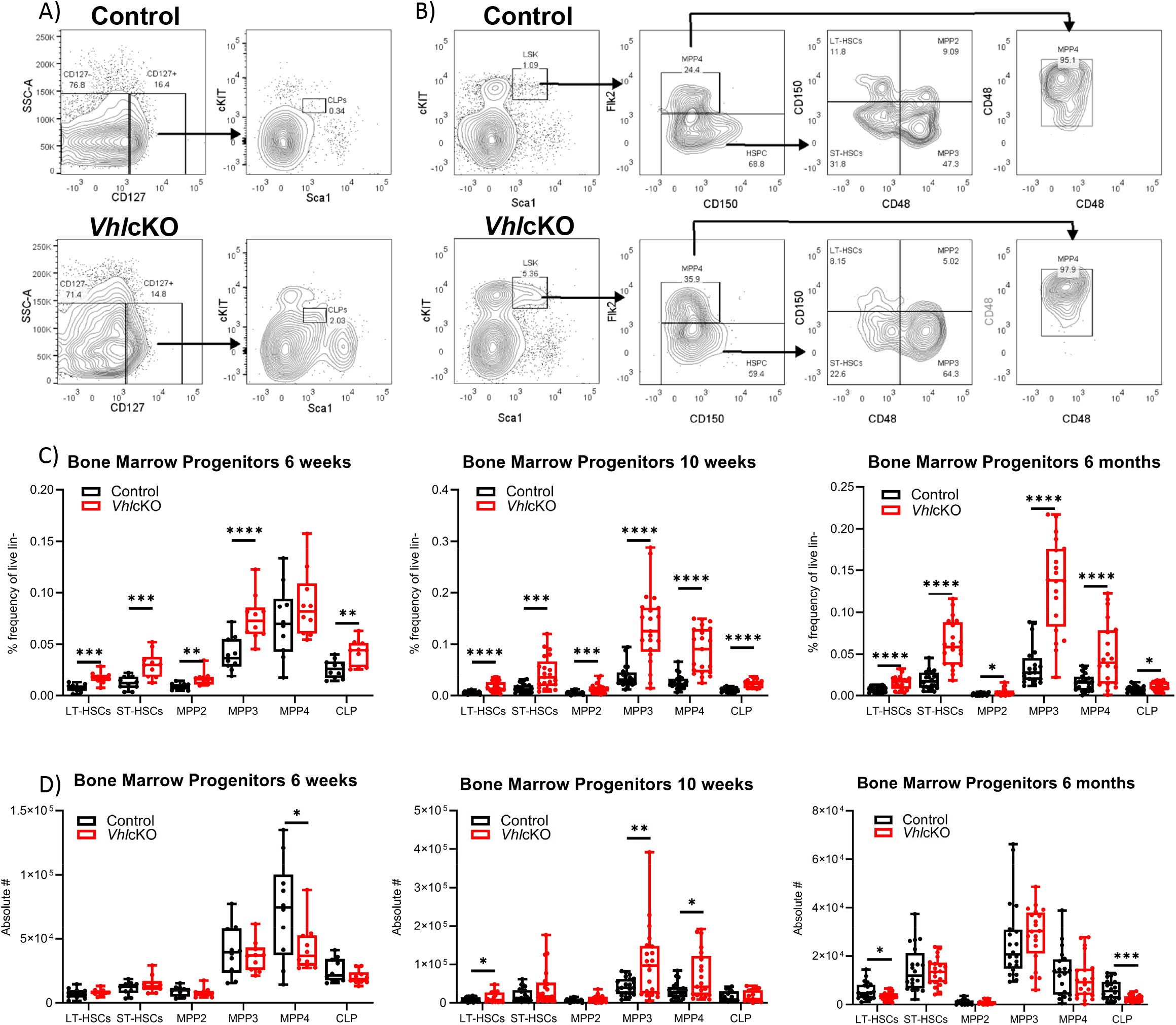
Altered B cell development in WT→*Vhl*cKO hematopoietic chimeras, demonstrating a cell-extrinsic effect of *Vhl*-deleted osteolineage cells on B cell development. A) Representative FACS plots of common lymphoid progenitors (CD127+ Sca1^int^ cKit^int^); B) representative FACS plots of hematopoietic progenitors in the bone marrow of controls (top) and *Vhl*cKOs (bottom); C) frequency and D) absolute number of hematopoietic progenitors in the bone marrow in 6-weeks-old (left), 10-weeks-old (middle) and 6-months-old (right) mice. p<0.05*, p<0.01**, p<0.001***, p<0.0001**** two-tailed Student’s t-test.

### *Vhl* deletion in *Dmp1*-expressing cells dysregulates B cell development in the BM

To further explore the effects of *Vhl* deletion in OBs and OCYs on B cell development and to identify at which stage B cell development was stunted in the BM, we determined the frequencies of Hardy Fractions A-F (**Figure 3A**, **Supplemental Figure 2**) using flow cytometry (1, 51). A decrease in frequencies of Fractions B-C and Fraction F was observed as early as 6-weeks-old, and across Fractions B through Fractions F was observed at 10-weeks-old and 6-months-old (**Figure 3B**). An overall decrease in the absolute numbers of B cells across all developmental stages was observed at all three ages (**Figure 3C**). *Vhl*cKO mice regardless of age displayed increased CLPs, retained normal frequency of Fraction A, whereas later Fractions were all decreased (**Figure 3B**). These results indicate an incomplete block in B cell development that starts at Fractions B-C in *Vhl*cKO mice.

**Figure 3.**
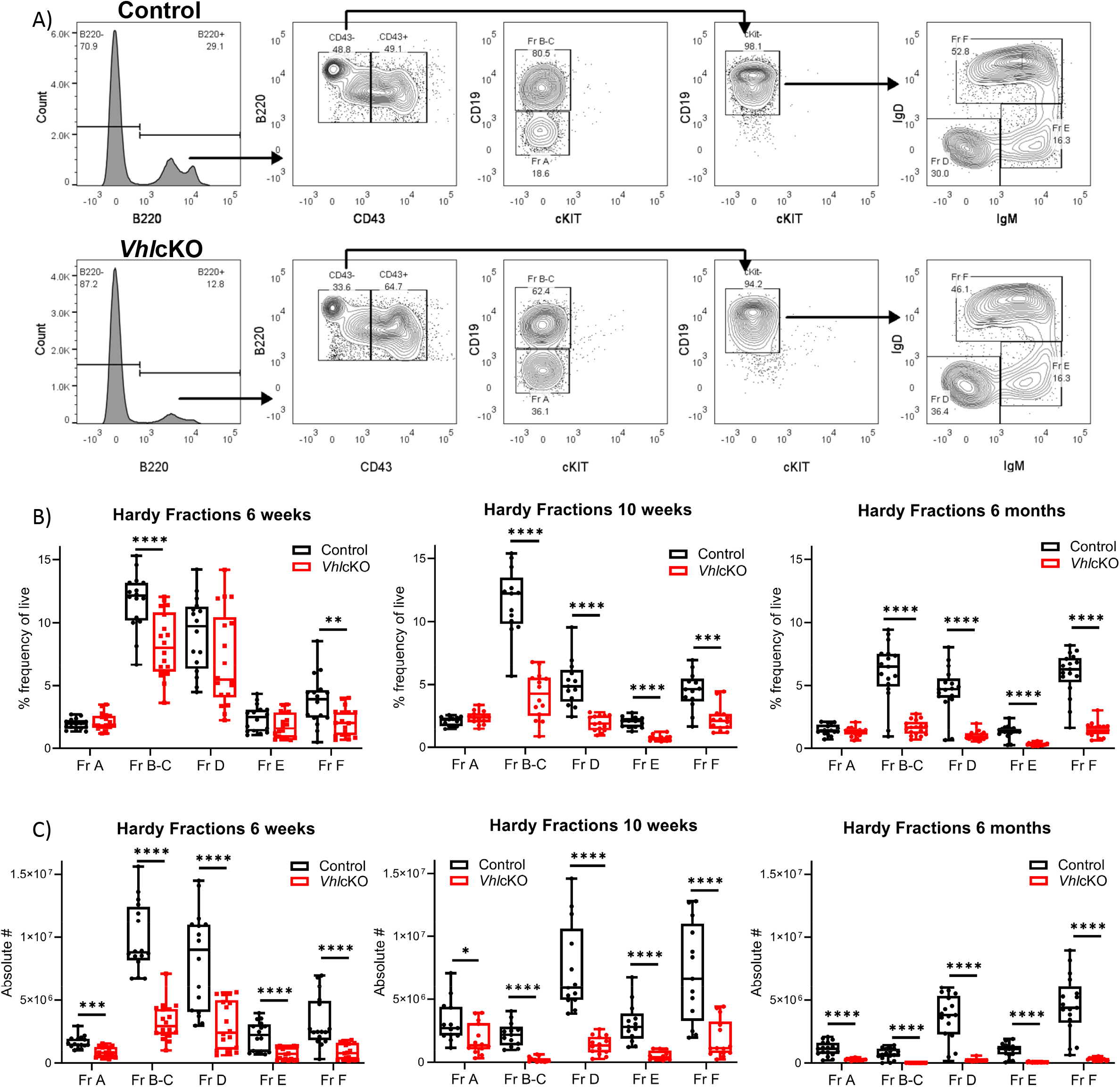
Dysregulated B cell development in *Vhl*cKO mice bone marrow. A) Representative FACS plot of B cell development in the BM control (top) and *Vhl*cKO (bottom); B) B cell frequency in 6-week-old (left), 10-week-old (middle), and 6-month-old (right) mice; C) absolute cell numbers in 6-week-old (left), 10-week-old (middle), and 6-month-old (right). p<0.05*, p<0.01**, p<0.001***, p<0.0001**** two-tailed Student’s t-test.

### Reciprocal bone marrow transplantation studies reveal a cell-extrinsic effect of the *Vhl*cKO microenvironment on B cell development

We expected the cause of the B cell defect to lie within the non-hematopoietic cells, since *Dmp1* is not expressed in hematopoietic cells. To definitively determine if the effects of *Vhl* deficiency on B lymphopoiesis were due to changes in the non-hematopoietic microenvironment within the bone, we performed whole BM transplants from WT (CD45.1+) donors into lethally irradiated *Vhl*cKO (CD45.2+) recipients (WT→*Vhl*cKO chimeras (**Figure 4A**)). Control WT (CD45.1+)→WT (CD45.2+) chimeras were also prepared. Donor hematopoietic chimerism was similar in controls and chimeras (**Figure 4B**). Analysis 16 weeks post-transplant showed a significant reduction in BM cellularity (**Figure 4C**) and an increase in granulocytes and monocytes, and a decrease in B cells in the WT→ *Vhl*cKO mice (**Figure 4D**). Analysis of B cells revealed a decrease at Fractions A through Fraction F in both frequency and absolute numbers (**Figure 4E, F**), similar to that observed in non-transplanted *Vhl*cKO mice (**Figure 3**). In contrast, overall hematopoiesis, including B cell development, was normal in the *Vhl*cKO→WT chimeras (**Supplemental Figure 3).** Since *Vhl* deletion in B cells can affect their function (52, 53), we confirmed that *Vhl* remained intact and was not erroneously deleted in B cells in our *Vhl*cKO mice (**Supplemental Figure 4)**. These results confirm a cell-extrinsic effect of the non-hematopoietic *Vhl*cKO BM microenvironment on hematopoiesis.

**Figure 4.**
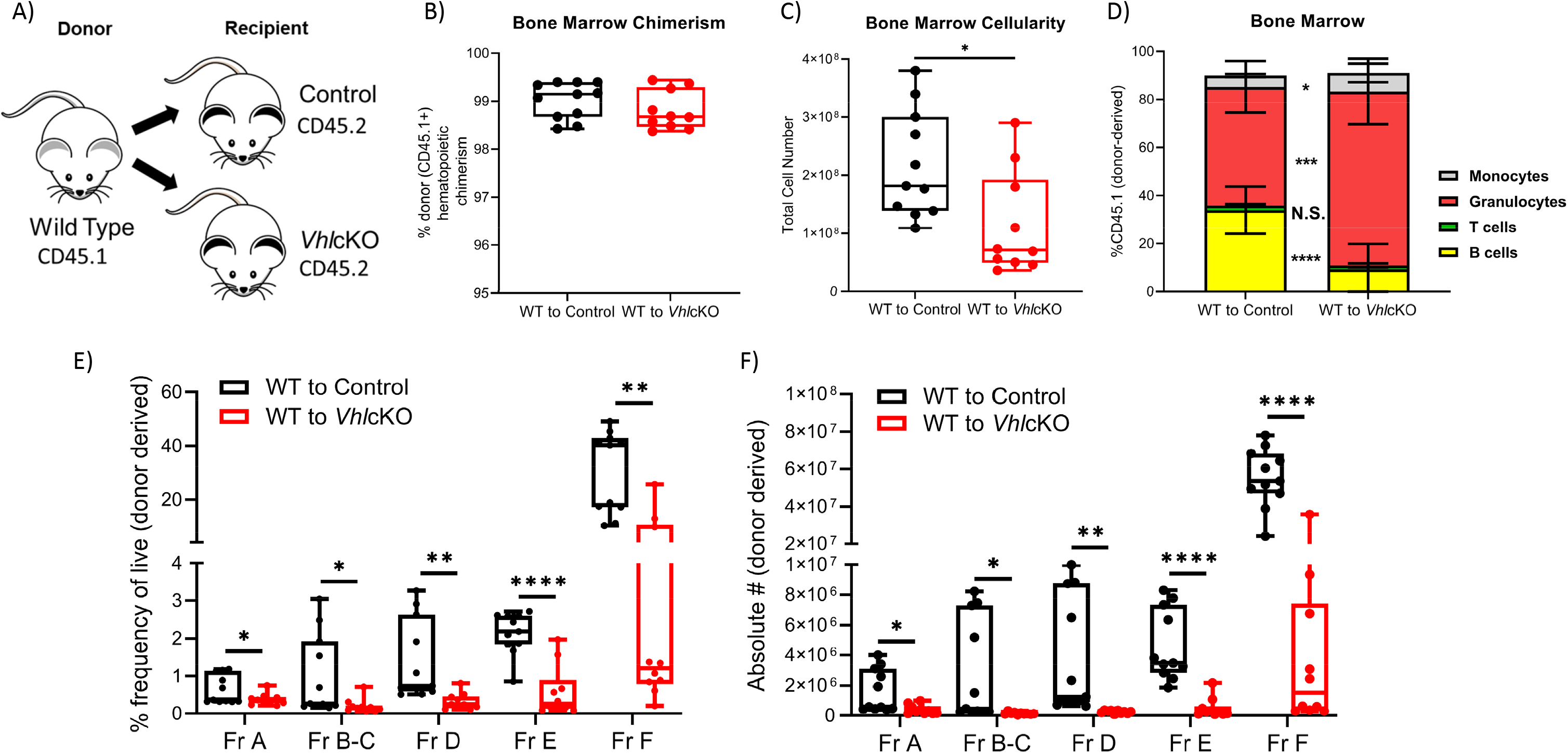
Altered B cell development in WT→*Vhl*cKO hematopoietic chimeras demonstrates a cell-extrinsic effect of the *Vhl*cKO bone on B cell development. Mice were transplanted at 10 weeks of age and were analyzed 16 weeks post-transplantation. A) experimental scheme; B) donor (CD45.1) chimerism; C) bone marrow cellularity; D) frequency of lineage cells in bone marrow; E) B cell frequency and F) absolute number of B cell developmental stages at 16 weeks post-transplant. p<0.05*, p<0.01**, p<0.001***, p<0.0001**** two-tailed Student’s t-test.

### *Vhl*cKO mice display patterns of reduced B cell proliferation and increased B cell apoptosis in the BM

We hypothesized that the observed reduction of B cells was due to increased apoptosis and diminished B cell proliferation in the BM. To test this, B cells were stained with Annexin V and 7AAD to identify cells that were live, in early stage apoptosis or late stage apoptosis (**Figure 5A**, **left panels**). Normally, apoptosis is the most extensive in Fraction A (pre-pro-B cells) amongst the B cell fractions (54). The frequencies of *Vhl*cKO Fraction A cells in live, early and late apoptosis stages was comparable to controls at 6- and 10-weeks-old. However, at 6-months old, Fraction A comprised fewer live cells and more late apoptotic cells compared to controls. Apoptosis in Fraction B-C in *Vhl*cKOs was similar to controls at 6-weeks-old, but then displayed an increase in late apoptosis at 10-weeks-old and 6-months-old (**Figure 5B**). A reduction of live Fraction B-C cells was also observed at 6-months old (**Figure 5B**). No differences in the stages of apoptosis were observed between controls and *Vhl*cKOs for Fractions D, E and F (**Supplemental Figure 5).**

**Figure 5.**
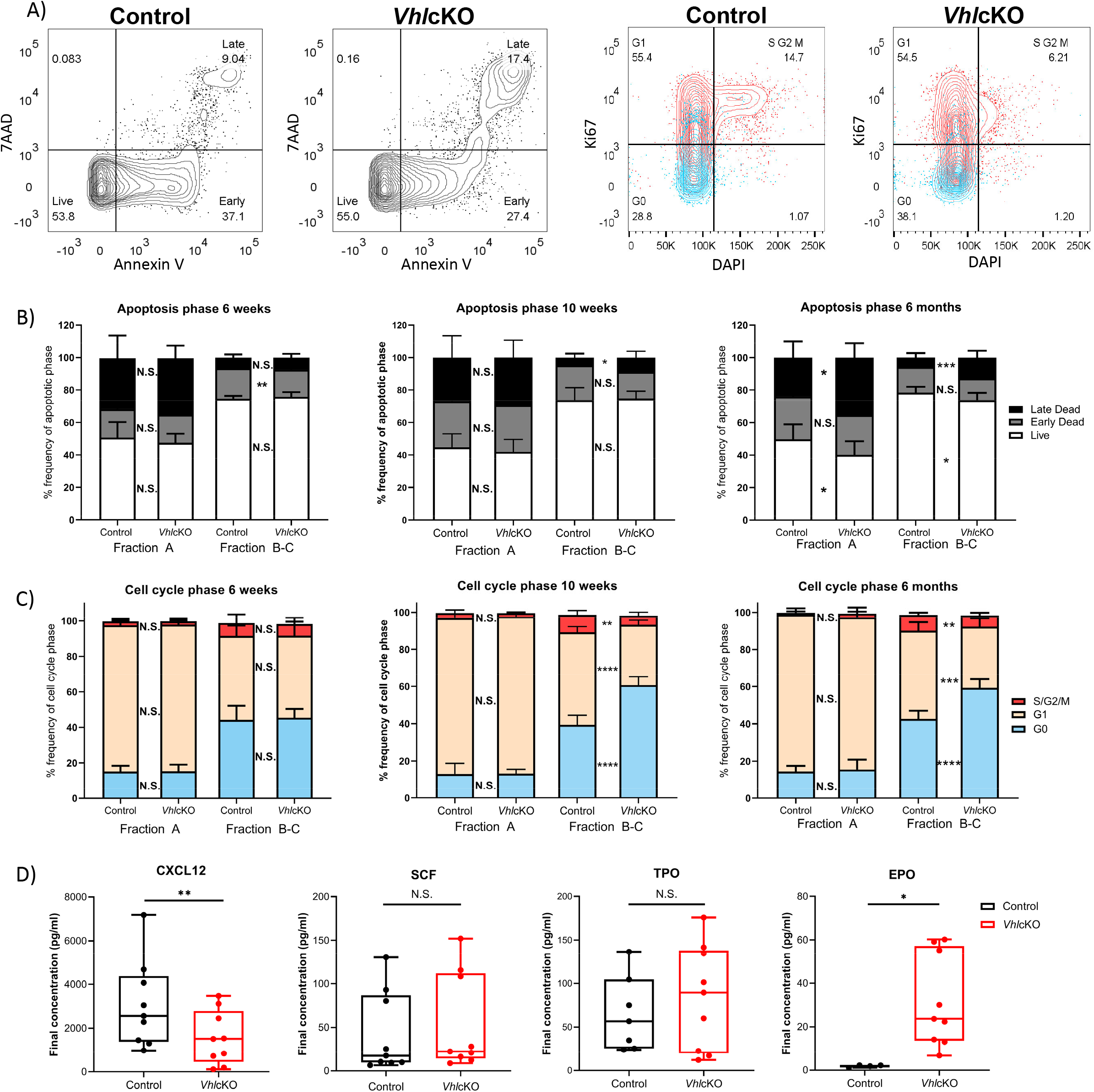
*Vhl*cKO mice display increase apoptosis and reduced cell proliferation during early B cell development. A) Representative FACS plots of apoptotic phases in B220+ cells (left) and cell cycle phases in B220+ cells (red:CD43+ blue: CD43-) (right); B) frequency of apoptotic phases in Fractions A and B-C in 6-weeks-old, 10-weeks-old and 6-month-old mice; C) frequency of cells in each cell cycle phase within Fractions A and B-C at 6-weeks-old, 10-weeks-old old and 6-month-old mice; (D) CXCL12, SCF, TPO, and EPO cytokine level measurements in bone marrow serum of control or *Vhl*cKO mice. p<0.05*, p<0.01**, p<0.001***, p<0.0001**** two-tailed Student’s t-test.

B cell development leads to the assembly and signaling of the B cell antigen receptor (BCR). CD43+ Fraction A-C (pre-pro-B and pro-B cells) normally have higher proliferation rates compared to CD43-Fraction D-E (Pre-B cells and immature B cells) (5, 55). Proliferation is halted at Fraction D (small pre-B cell) to allow light (L) chain gene rearrangement, subsequently expressing a complete IgM surface molecule (Fraction E) (5, 56). Cell cycle analysis was performed using Ki67 and DAPI staining (**Figure 5A**, **right panels**). At 6-weeks-old, there were no differences in the distribution of cells in G0 (quiescent), G1, or S2/G2/M phases between *Vhl*cKO and control mice. However, at 10-weeks-old and 6-months-old, Fractions B-C contained a reduced percentage of cells in G1 and S/G2/M cell cycle phases, and an increased percentage in G0 (**Figure 5C**). This indicates a reduced ability of early B cell progenitors to proliferate in a *Vhl*-deficient microenvironment. No difference in proliferation of Fractions D-F was observed at any age examined, with the exception of a slight (yet statistically significant) reduction of the *Vhl*cKO Fraction F cells in G0 and increase in G1 at 6-months-old (**Supplemental Figure 6**).

B cell development at each stage requires specific signaling molecules from a variety of niche cells (5, 57). To examine for changes in the distribution of niche cells in *Vhl*cKO bones, control and *Vhl*cKO long bones were digested and stained for stromal cell populations by flow cytometry (**Supplemental Figure 7)** (58, 59). Using this approach, the *Vhl*cKO mice displayed reduced frequency of ECs, while MSCs, and OBs remained similar to controls. To further explore the dysregulated niche, BM serum was analyzed for levels of CXCL12 and SCF, which are critical for B cell development (1, 24, 28, 29). CXCL12 levels were reduced in the *Vhl*cKO BM serum, while SCF levels were unaffected (**Figure 5D**). This suggested that increased apoptosis and reduced proliferation of Fraction B-C cells are caused by reduced CXCL12 levels and lack of ECs in the *Vhl*cKO BM.

### Evidence for elevated oxygen levels in local niches in the *Vhl*cKO bone marrow

Hypoxic niches in the BM microenvironment are crucial for hematopoiesis development. Dynamic regulation of HIF-1α levels are required for normal B cell development such that HIF activity is high in pro-B and pre-B cells and decreases in the immature B cell stage in the BM (60). In wild type mice, studies using the hypoxic marker pimonidazole (PIM) revealed that HSCs in the BM stain positively with PIM, indicating a hypoxic niche (61). To evaluate hypoxia in distinct B cell developmental stages, *Vhl*cKO and control mice were injected with PBS or 120 mg/kg PIM. PIM staining of LSKs in the BM stained positively for PIM, as previously reported by other groups (**Figure 6A**). CD45+ B220+ cells (which include all Hardy Fractions, in addition to other hematopoietic progenitors, natural killer cells, dendritic cells and T cells (62–66)) displayed positive, yet less intense staining with PIM in both control or *Vhl*cKO mice (**Figure 6A**). Normalization of PIM staining on LSK HSCs and CD45+ B220+ cells in four *Vhl*cKO mice to the mean staining in controls showed no statistically significant differences (**Figure 6B**). However, examination of PIM staining in distinct Hardy Fractions revealed that in general, the early B cell progenitors (Fraction A) stain with PIM at a higher level than the mature B cells (Fraction F). Notably, *Vhl*cKO mice displayed diminished intensity of PIM staining compared to control mice across all B cell fractions (**Figure 6C**, **Table 2**). This reveals that similar to LSKs, Fraction A cells might reside in a hypoxic niche, and as they mature they move away to a less hypoxic niche. Moreover, these data indicate that B cells in the *Vhl*cKO BM may experience higher oxygen levels as compared to control mice in their local niches.

**Figure 6.**
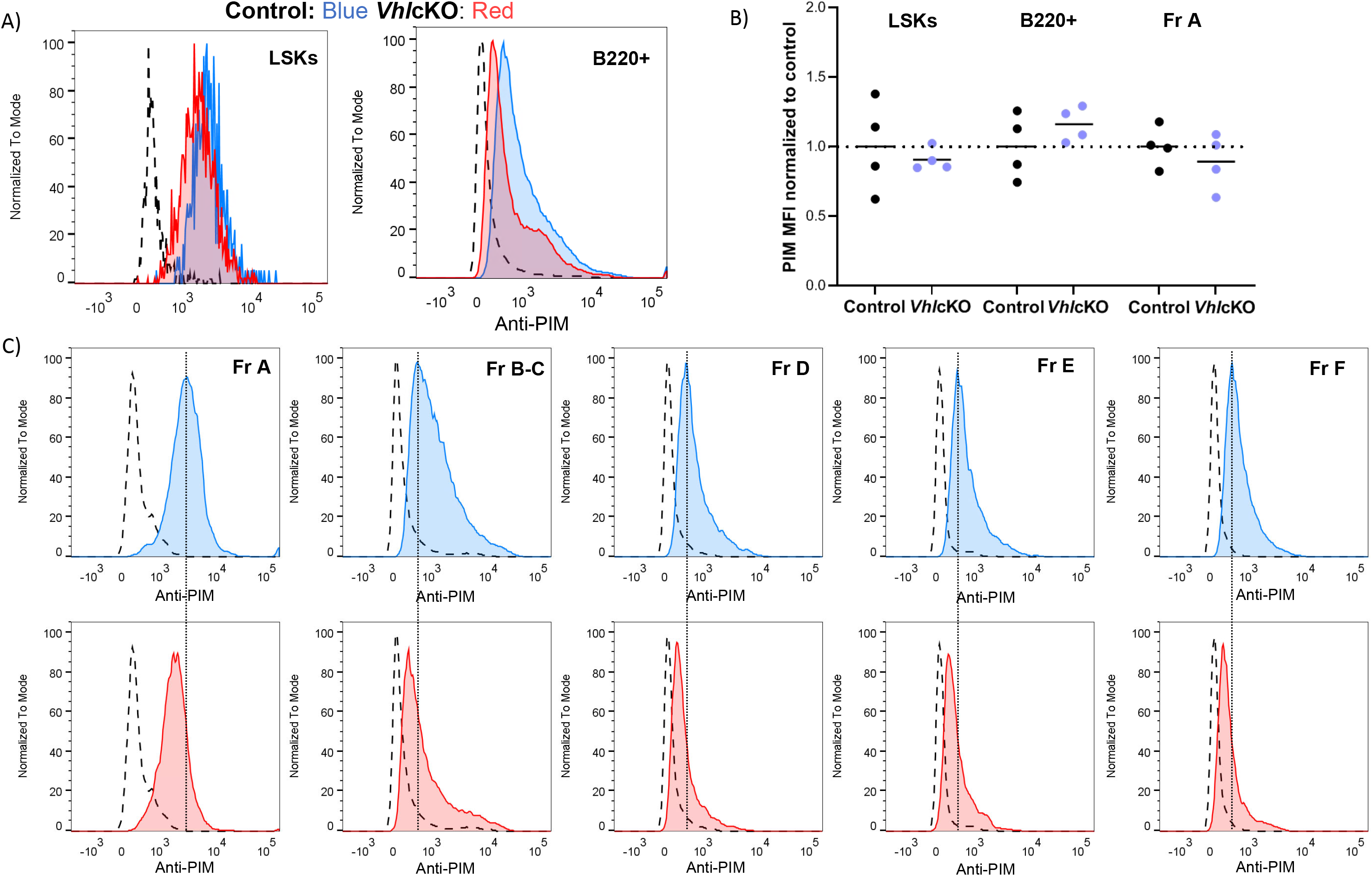
Hypoxia cell marker pimonidazole indicates difference amongst B cell fractions in control and VhlcKO mice. *Vhl*cKO and control mice were injected with PBS or 120 mg/kg pimonidazole (PIM). PIM staining of A) LSKs in the BM and CD45+ B220+ cells; dashed line: isotype control; blue line: anti-PIM staining in control (WT); red: anti-PIM staining in *Vhl*cKO; B) summary of PIM staining in four *Vhl*cKOs (purple dots) normalized to the mean fluorescence intensity (MFI) in WT controls (black); results from 2 independent experiments are shown; C) representative anti-PIM staining plots of individual B cell Fractions A-F from a control (top) and a *Vhl*cKO mouse (bottom). The thin dashed line crossing over the control and *Vhl*cKO plots is intended to help visualize the shift in PIM fluorescence intensity in the *Vhl*cKO samples.

**Table 2.** Mode Fluorescence Intensity of PIM staining on B cell fractions in control and *Vhl* cKO.

| **Genotype** | **Treatment** | **MFI (mode)** |  |  |  |  |
| --- | --- | --- | --- | --- | --- | --- |
|  |  | **Fr A** | **Fr B-C** | **Fr D** | **Fr E** | **Fr F** |
| *Vhl* cKO | Isotype | 336 | 125 | 146 | 187 | 146 |
| *Vhl* cKO | Isotype | 358 | 166 | 166 | 166 | 208 |
| Control | PIM | 3561 | 742 | 588 | 663 | 613 |
| Control | PIM | 3234 | 769 | 663 | 663 | 689 |
| Control | PIM | 2292 | 493 | 470 | 470 | 402 |
| Control | PIM | 3926 | 769 | 715 | 663 | 663 |
| *Vhl* cKO | PIM | 2514 | 493 | 402 | 336 | 424 |
| *Vhl* cKO | PIM | 2593 | 613 | 516 | 540 | 588 |
| *Vhl* cKO | PIM | 3926 | 689 | 570 | 663 | 564 |
| *Vhl* cKO | PIM | 3561 | 588 | 516 | 493 | 516 |

### Increased bone marrow blood vessel diameter and density in *Vhl*cKO microenvironments

To more precisely examine the changes in the microenvironment of *Vhl*cKO mice, we imaged femurs that were shaved to remove cortical bone (for analysis of the metaphysis) or optically cleared with a modified uDISCO protocol (for analysis of the fully intact diaphysis) **(Supplemental Videos 1-2)** (43). We measured the vessel diameter and frequency in the cleared long bones and found that regardless of their position in the BM, blood vessels in *Vhl*cKO mice were significantly larger than the control group **(Figure 7A-C)** while generally no difference was observed in the vessel frequency (**Supplemental Figure 8A)**. Bone and BM vessel density measurements in both the metaphysis and diaphysis revealed that in *Vhl*cKO, blood vessels occupy a larger volume than controls **(Figure 7D-F**, **Supplemental Figure 8B-C)**. Furthermore, we observed an apparent decrease in endosteal lining arterioles in the diaphysis of 6-month-old *Vhlc*KO femurs compared to controls (**Supplemental Figure 8D**). Taken together, these data reveal a striking alteration in the overall architecture of the BM vascular network in *Vhlc*KO mice.

**Figure 7.**
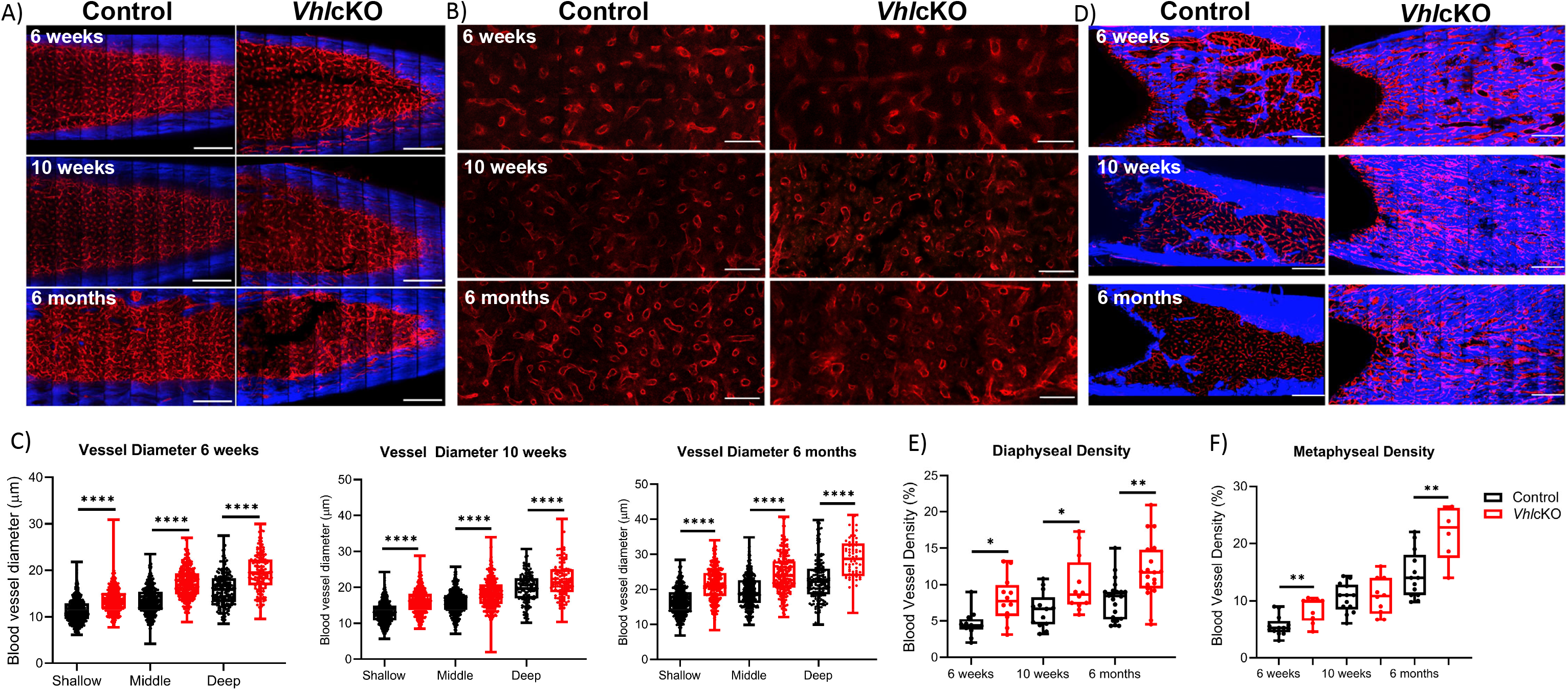
Ex vivo two-photon imaging of long bones in *Vhl*cKO and controls. A) Representative macroscopic images of the femur diaphyseal BM (scale bars: ~200 μm), B) magnified z-stacks (scale bars: ~100 μm), and C) statistical analysis after uDISCO clearing show an increase in the *Vhl*cKO vascular diameter relative to the controls; D) *ex vivo* images of femur metaphyseal BM after max intensity projection reveal bone replacement and vascular alteration in *Vhl*cKO; E-F) quantification of the metaphyseal and diaphyseal vascular density (scale bars: ~200 μm). Red: blood vessels (labeled with Alexa647 conjugated antibodies against CD31, CD144, and Sca-1), Blue: bone (SHG: Second harmonic generation). *p<0.05, **p<0.01,***p<0.001, ****p<0.0001, two-tailed Student’s t-test.

### *Vhl*cKO bone marrow blood vessels display increased permeability

While it has been shown that the bone and vascular system undergoes significant remodeling in *Vhl*cKO mice, there has been a lack of information regarding potential functional changes to BM blood vessels. To examine changes to the BM vasculature system which could negatively impact B cell development, we sought to quantify changes to the vascular permeability, leakage and blood flow velocity via intravital two-photon microscopy of the calvaria. Vessel permeability reflects the rate at which small molecules exit blood vessels and fill the surrounding perivascular space, whereas leakage is the ratio of fluorescent dye in the perivascular space and vascular lumen after reaching equilibrium. Blood vessel leakage and permeability was calculated by administering Rhodamine B Dextran (70kDa) via a retro-orbital injection. We found that *Vhl*cKO mice displayed greater vascular leakage overall, and that vascular leakage increased in both control and *Vhl*cKO mice with age (**Figure 8A-B**, **Supplemental Videos 3-8**). Similarly, we observed an increase in vascular permeability in *Vhl*cKO mice, which significantly increased with age (**Figure 8C**, **Supplemental Video 9-10**). We observed a decrease in blood flow velocity in *Vhl*cKO mice compared to controls for 6-week-old and 10-week-old mice (**Figure 8D**). Lastly, we observed an age-related reduction in blood flow in both *Vhl*cKO and control mice (**Figure 8D**), which is consistent with previously published changes in BM vascular flow rate with age (67).

**Figure 8.**
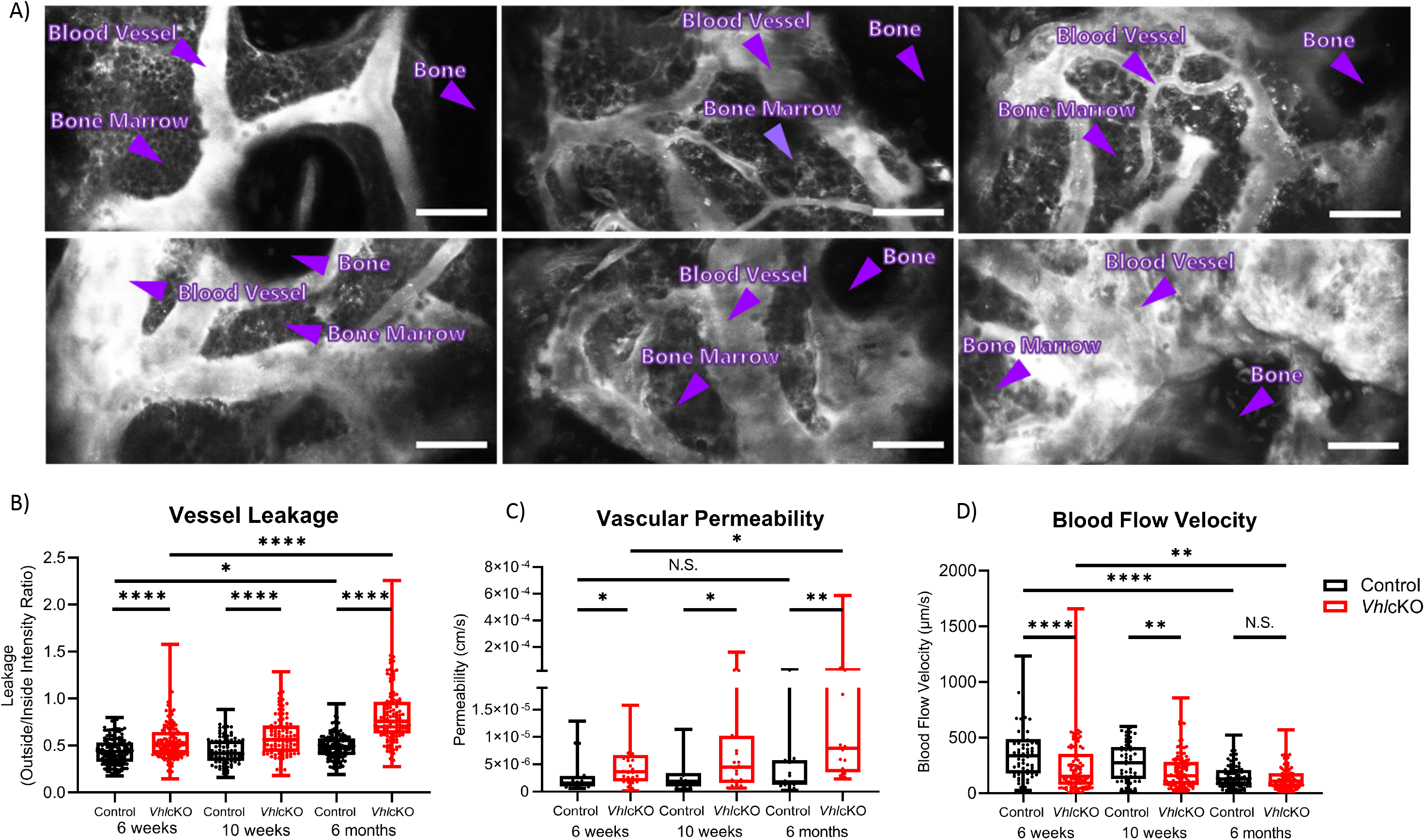
Disruption in blood-bone marrow barrier revealed by intravital microscopy. Blood vessel microenvironment comparisons of control and *Vhl*cKO mice at 6-week (n=4), 10-week (n=4) and 6-month (n=5) timepoints. A) Representative contrast adjusted max intensity projections of the calvarial BM in control and *Vhl*cKO mice by age; White: blood vessel (Rhodamine B Dextran, 70 kDa); scale bar: 50 μm; quantification of calvarial BM B) blood vessel leakage, C) vascular permeability, and D) blood flow velocity. *p<0.05, **p<0.01, ***p<0.001, ****p<0.0001, Mann-Whitney test.

## DISCUSSION

VHL plays an important role regulating HIF expression, and disruption of *Vhl* in bone cells leads to improper bone homeostasis (7, 8, 19, 37). *Vhl* deletion in osteochondral progenitor cells and osteocalcin-positive OBs leads to an increase in bone mass through an increase in OB number (7, 19). Furthermore, disrupting VHL in OBs induces expression of β-catenin, revealing that VHL/HIF pathway promotes bone formation through the Wnt pathway (7, 20, 21). Altogether, these studies of *Vhl* deletion in osteolineage cells have not examined the cell-extrinsic effects of these changes on the immune cells residing in the BM. In this report, we establish that deletion of *Vhl* in late OBs and OCYs results in cell-extrinsic changes that does not support full development and survival of B cells.

BM stromal cells include a variety of non-hematopoietic cells, such MSCs and OBs, which are the precursors of mature mineralized OCYs, and ECs. All of these cells support B cell development (1, 28, 31, 32). In our study, using flow cytometry after bone digest, we found that the distribution of the stromal cell populations was not affected in the *Vhl*cKO, with the exception of CD31+ ECs, which were reduced. This was surprising due to the increase in vessel volume and number that was observed in the ex vivo imaging of *Vhl*cKO long bones. The differences in the results may reflect a limitation in the bone digestion protocol that is used for analyzing BMSCs by flow cytometry. It is widely accepted that with this protocol, hematopoietic cells, likely from the bone marrow, are still evident after rinsing of the bone chips and collagenase digestion. Therefore, we cannot determine if the ECs examined are residual BM ECs or resident bone ECs. In addition, the increased density of *Vhl*cKO bone makes it challenging to digest completely and all of the ECs from the bone may not be released. Given this caveat, we believe that our imaging results are more accurate than the flow cytometry, and the imaging results suggest that there is an increase in bone ECs, which is consistent with previous studies in *Osx-*Cre;*Vhl^fl/fl^* mice where endomucin staining showed that *Vhl* deletion increased bone vasculature with dilated blood vessels (37). We also observed larger vessels in the BM across all ages and an increase in BM blood volume. These changes, along with the observed decrease in endosteal arterioles in the long bones, suggests that oxygenation of the *Vhl*cKO marrow may be higher than normal, which may play a role in dysregulation of B cell development. Future studies will be needed to clarify this and to identify other changes in specific types or locations of blood vessels in the *Vhl*cKO model.

Given the connection between *Vhl* and hypoxia response, it was interesting that EPO levels were high in the BM serum of the *Vhl*cKO mice. This has been confirmed in other studies, where deletion of *Vhl* at the mature OB stage using the *Osx*-Cre and *Ocn-*Cre (targeting osteoprogenitors) (8, 19, 20) and at the late OB/OCY stage using *Dmp*-Cre increased bone mass and angiogenesis, likely through HIF1α-regulated expression of VEGF and EPO. If the elevated EPO levels directly affect B cell development in the *Vhl*cKO BM has not yet been verified. However, it has been reported that ECs in the BM suppress levels of CXCL12 expression in response to increased EPO levels (68). We also observed decreased CXCL12 in the BM sera of *Vhl*cKO mice. CXCL12 is required for proper development and retention of B cells in the BM (32, 69). This suggests that altered vascular components in the *Vhl*cKO bone and BM microenvironments impair B cell development possibly through the effects of EPO on EC function.

Permeability of the BM vasculature in the *Vhl*cKO mice was also compromised. We found an increased vascular leakage and permeability in the *Vhl*cKO BM compared to controls regardless of age. In addition, vascular permeability appeared to increase with age, with the highest vascular permeability and leakage being observed in 6-month-old *Vhl*cKO mice when compared with 6-week-old mice. Interestingly, it was observed that vascular blood flow velocity decreased in 6-week-old and 10-week-old *Vhl*cKO mice but was not affected in 6-month-old *Vhl*cKO mice. An increase in blood flow velocity would normally explain an increase in permeability and leakage, but that is not evident in our data. Instead, the more likely explanation is that the blood-bone marrow barrier is compromised, increasing the exposure of the BM to plasma components.

Deletion of *Vhl* in B cells stabilizes *Hif1α* levels and affects mature B cell function by impairing cell proliferation, antibody class-switching, generation of high affinity antibodies, antibody responses, and impairs metabolic balance essential for naive B cell survival and development (52, 53, 70). Dynamic regulation of HIF-1α levels was also found to be a crucial step in B cell development in the BM (60). Burrows et al. found decreased *Hif*1α activity at the immature B cell stage in the BM and that HIF-1α suppression was required for normal B cell development (60). This dynamic regulation of HIF-1α activity during B cell development is consistent with our results, which revealed that Fraction A cells stain highly with PIM, and PIM staining was reduced as B cell development progressed to Fraction F. Together, our findings and that of Burrows et al. suggest that the earliest B cell stages (e.g. pre-pro B, Fraction A) might prefer a more hypoxic niche compared to the later B cell stages. We also found that the *Vhl*cKO mice displayed lower intensity of PIM staining on B cells compared to controls, suggesting a hyperoxic state, which could also be deleterious for B cell development. Although *Vhl* is deleted in OBs and OCYs in our model, we cannot yet rule out if this deletion is artificially causing changes that would be found in a true hypoxic state through *Hif1* stabilization, when in fact the oxygenation of the BM of the *Vhl*cKO is not altered. In addition, PIM cannot provide true quantification of dissolved oxygen concentration in tissue. PIM adduct staining results could reflect inadequate oxygen supply to the BM, faulty rates of intracellular oxygen consumption, or both. Direct in vivo measurement of oxygen tension using two-photon phosphorescence lifetime microscopy would help answer this question (11).

HSCs were increased in previous studies that showed that HSCs can maintain cell cycle quiescence and function through regulation of HIF-1α levels (9). In our studies, we found that HSCs and upstream progenitors of B cells were increased in frequency. We found that the increase in progenitors is due to the inability to mature into B cells, but it remains unclear whether there is a shift from lymphoid to myeloid skewing. Some possible studies to sort out lineage bias would be through immune phenotyping analysis such as single cell sequencing to explore expression signatures (myeloid vs. lymphoid) to reveal lineage fate (49) or limiting dilution analysis and single cell transplantation as previously done by other groups to show the ratio of myeloid to lymphoid cell output (71–74).

The information generated in this study helps define the role of *Vhl* and altered bone homeostasis on immune cell development. Our results suggest the following working model of the interactions in the BM microenvironment that controls B cell development (**Figure 9**): *Vhl* in OCYs and late OBs play a significant role in the BM microenvironment, indirectly regulating B cell development through a decrease in CXCL12, an increase in EPO, increased vasculature and vascular permeability. However, oxygen tension in the niche of early stage B cell fractions is yet to be determined. Our results demonstrate the significant changes of the physical niche in *Vhl*cKO mice and their effects on B cell development. Whether the physical space, niche cells, or molecular signals all play a direct or indirect role on B cell development remains to be explored and defined, with the possibility that these events are completely independent of each other. The results of this work could contribute to the development of new therapies or new targets for exogenous CXCL12 and EPO antagonists, to preserve and improve bone marrow function during microenvironmental niche changes or stress.

**Figure 9.**
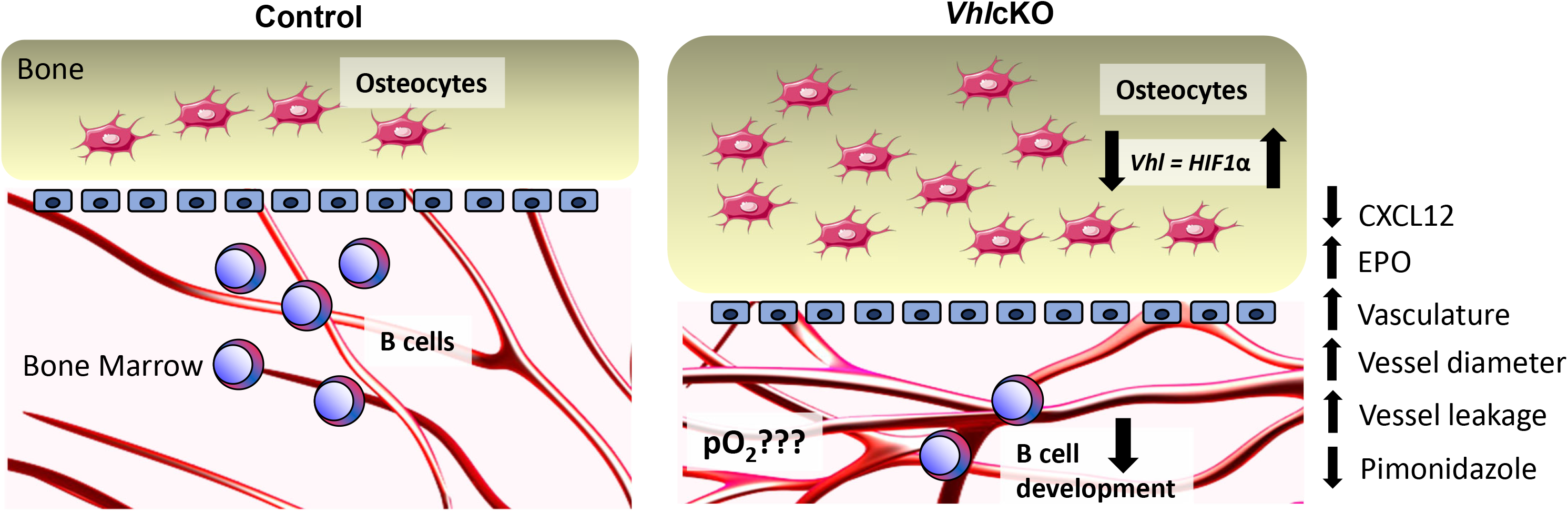
Working model describing the changes in the bone marrow microenvironment in *Vhlc*KO mice. Left panel: Schematic of healthy control bone marrow where VHL/HIF signaling is intact, the transition from osteoblasts to osteocytes is homeostatically balanced and interactions of developing B cells and stromal cells within their niches promotes their differentiation, maturation and proliferation. Right panel: Lack of *Vhl* in late osteoblasts and osteocytes has a severe effect on hematopoiesis in the bone marrow, changing the B cell niche and indirectly regulating B cell development through decrease of CXCL12, increase of EPO and changes to the BM microenvironment vasculature and permeability. Direct measurement of pO_2_ in the BM is necessary to determine if the BM oxygenation landscape is altered compared to controls.

## ACKNOWLEDGMENTS

We thank the staff of the Department of Animal Research Services (DARS) and the Stem Cell Instrumentation Foundry (SCIF) at UC Merced for the excellent animal care and technical support. We also acknowledge Hawa Padmore and William Pratcher for their early contributions to protocol setup. The authors also thank the staff of the Health Sciences Research Institute (HSRI) at UC Merced for their administrative support. This work was funded by the University of California, NIH grants R15 AI154245-01 (J.O.M. and J.A.S.) and NIH F31 AI154815 (B.C.), and through support of the NSF-CREST: Center for Cellular and Biomolecular Machines at the University of California, Merced (NSF-HRD-1547848; N.A., C.B., and J.A.S.).

