## Supplemental Materials for "Deletion of *Vhl* in *Dmp1*-expressing cells causes microenvironmental impairment of B cell lymphopoiesis"

### SUPPLEMENTAL FILES

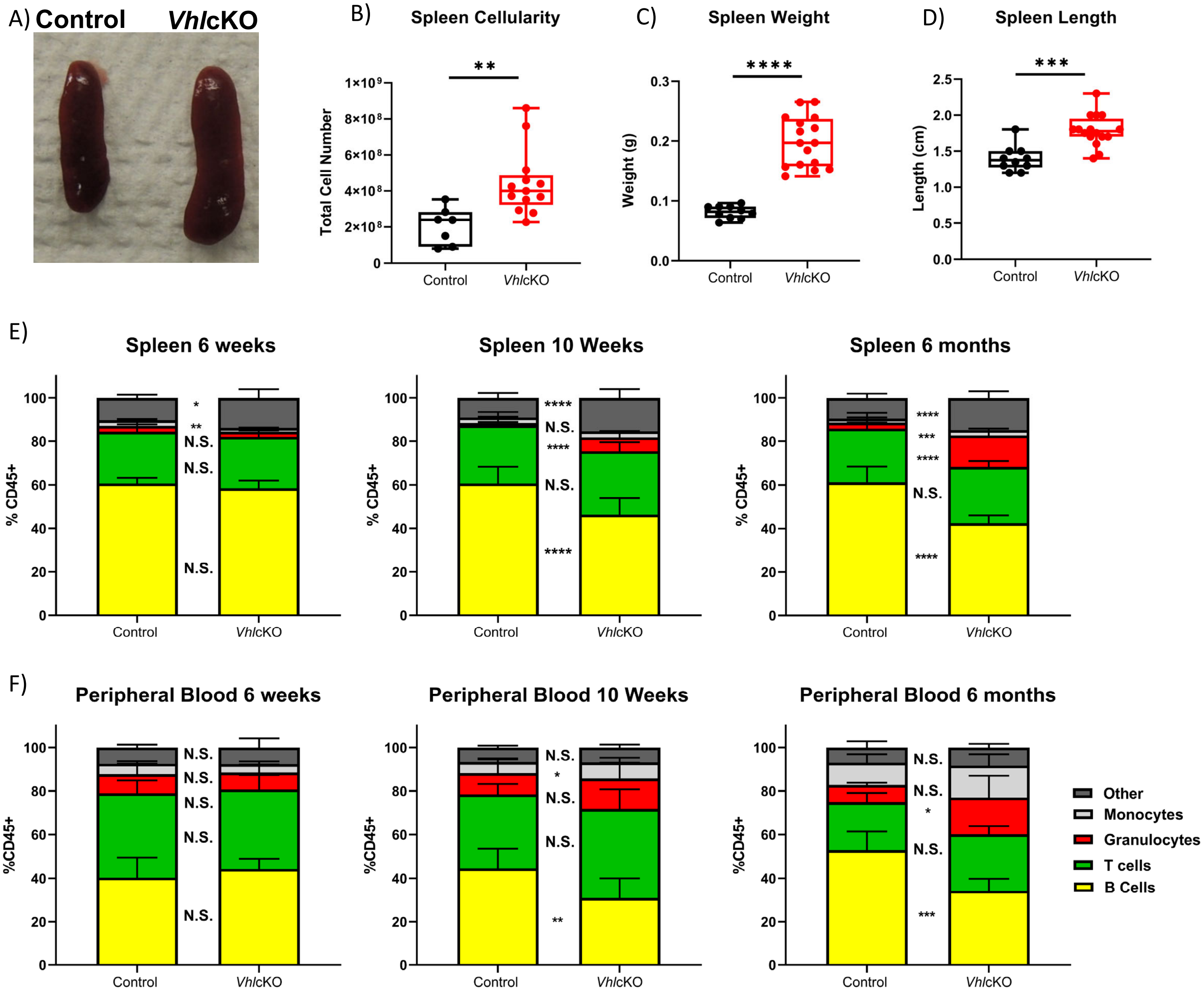


**Supplemental Figure 1. Analysis of hematopoietic lineages and cellularity in the spleen of *Vhlc*KO mice.** A) White light image of spleens, scale bar: 1cm; B) spleen cellularity; C) spleen weights; D) spleen length; E) spleen and F) peripheral blood lineage frequency at 6-weeks-old (left), 10-weeks-old (middle), and 6-months-old (right). *p<0.05, **p<0.01,***p<0.001, ****p<0.0001, two-tailed Student's t-test.

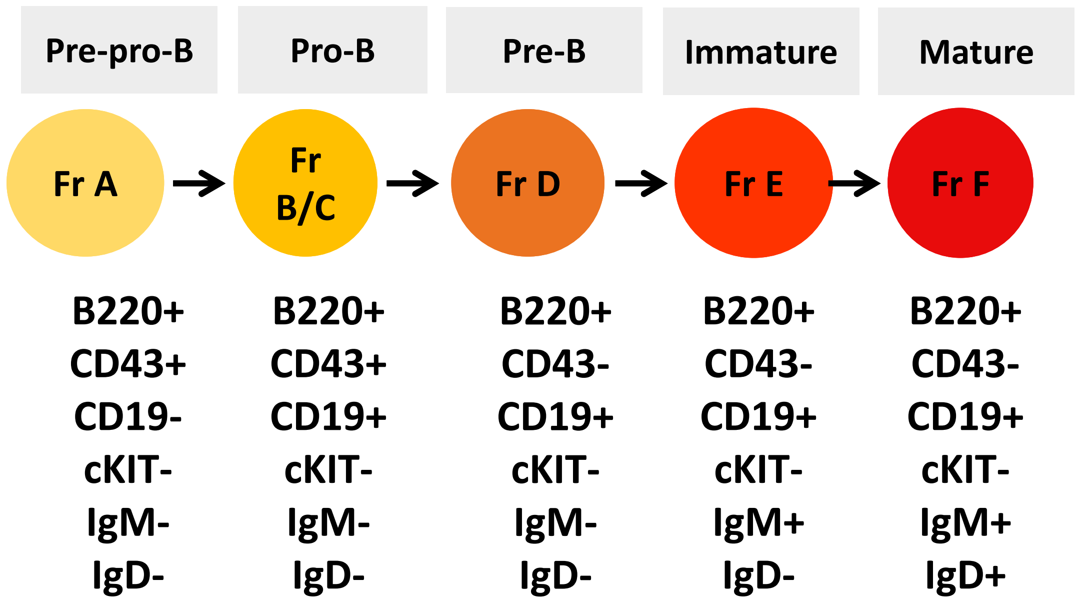


**Supplemental Figure 2. Antigen markers used to identify and distinguish B cell development stages.** Markers used for the analysis of B cell development at each developmental stage by flow cytometry. Hardy and Basel nomenclatures for B cell developmental stages are shown (51, 75).

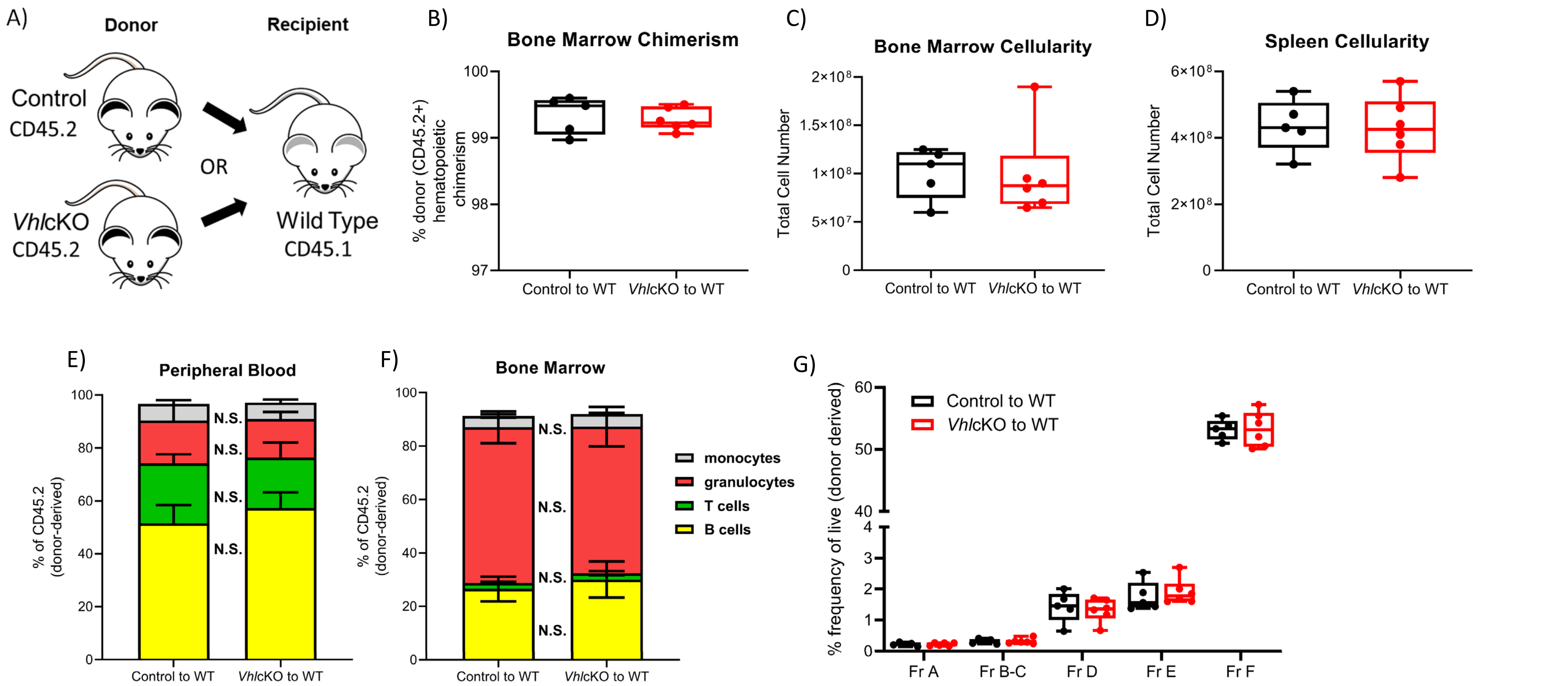


**Supplemental Figure 3. Evidence against cell-intrinsic effects of *Vhl* deletion on B cell development in *Vhl*cKO mice.** A) Experimental scheme. Mice were transplanted at 10 weeks of age and were 26 weeks old at time of analysis. B) donor (CD45.2+) chimerism; C) bone marrow cellularity; D) spleen cellularity; E) frequency of lineage cells in peripheral blood and in F) bone marrow; G) frequency of B cell developmental stages in chimeras 16 weeks post-transplant. p<0.05*, p<0.01**, p<0.001***, p<0.0001**** two-tailed Student's t-test.

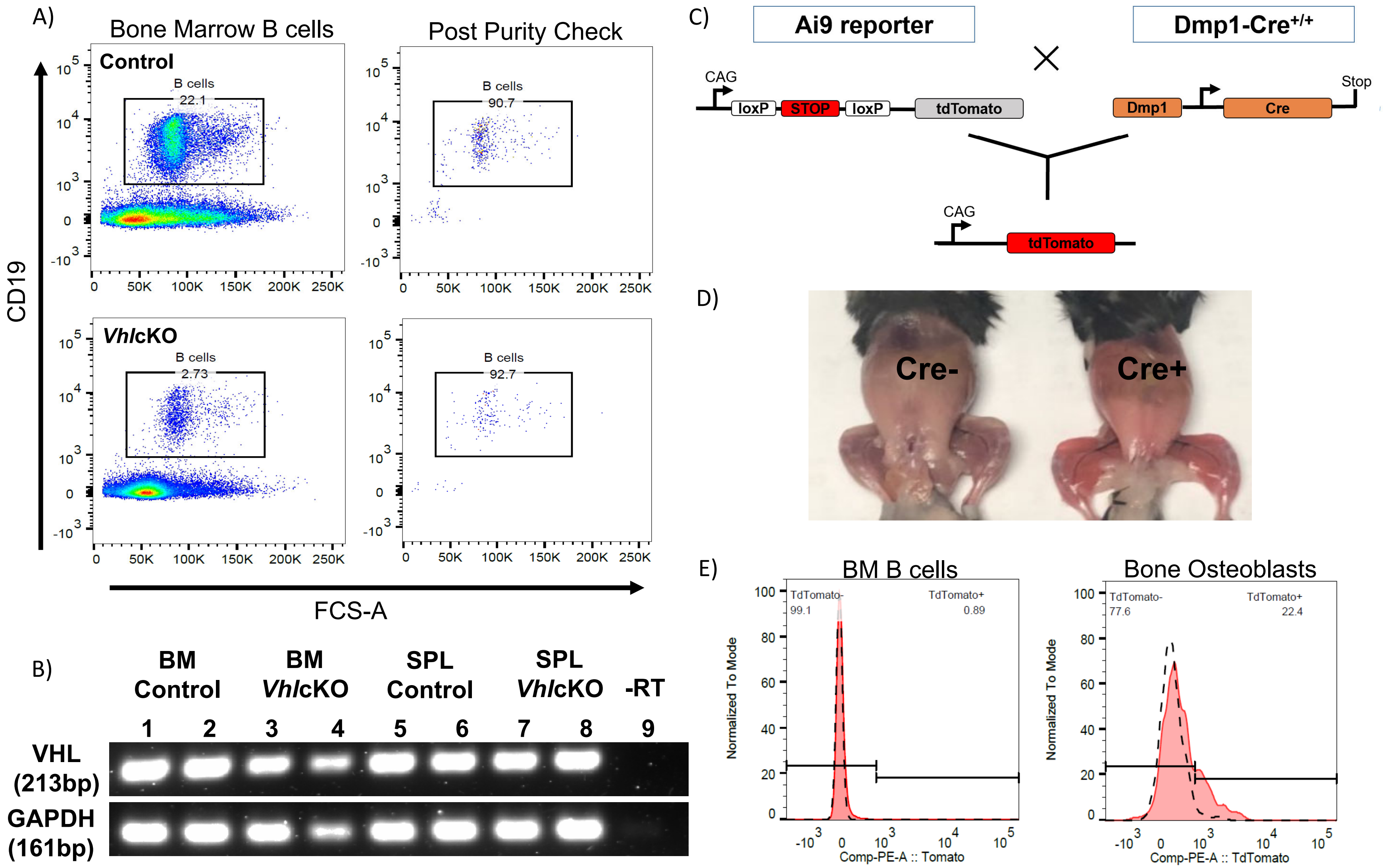


**Supplemental Figure 4. Validation of *Vhl* expression in B cells in *Vhl*cKO mice and expression of *Dmp1*-Cre using Ai9 reporter mice** A) Bone marrow CD19+ B lymphocyte percentages pre- and post- sorting from control and *Vhl*cKO mice and B) PCR validation of *Vhl* and *Gapdh* gene expression in DNA of sorted B cells (Live, CD19+), demonstrating *Vhl* is intact in B cells in the *Vhl*cKO; C) reporter cross of *Dmp1*-Cre mice with Ai9 (tdTomato) mice; D) tdTomato expression in *Dmp1*-Cre;Ai9 mice; E) flow cytometry measurement of tdTomato on BM B cells and osteoblasts (Lin-, CD45-, CD31-, Sca1-, CD51+) in *Dmp1*-Cre;Ai9 mice.

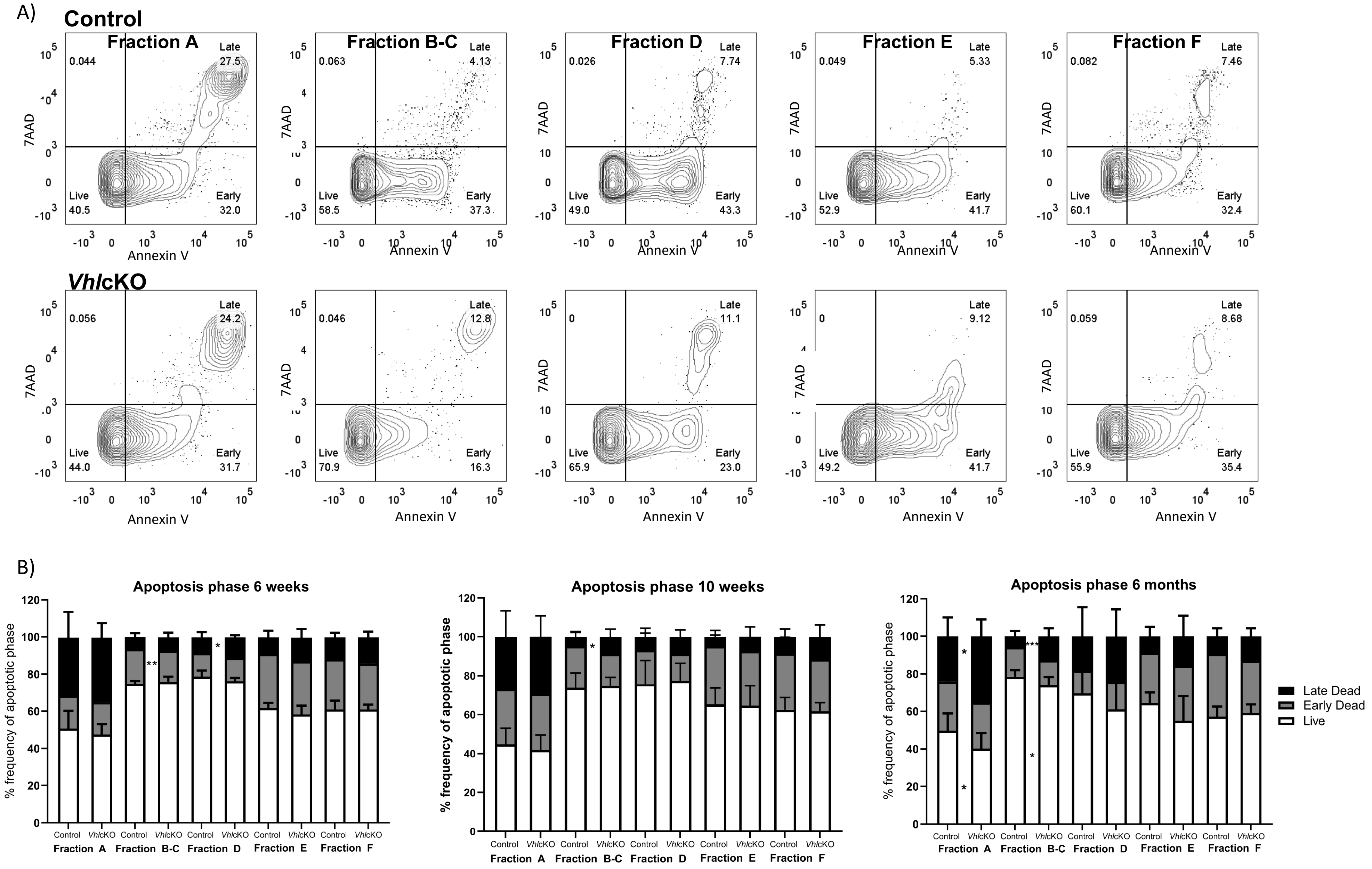


**Supplemental Figure 5. Apoptosis analysis of B cell Fractions A-F** A) Representative FACS plot of apoptotic phases in control (top) and *Vhl*cKO (bottom); B) frequency of apoptotic phase in Fractions A-F in 6-weeks-old, 10-weeks-old and 6-month-old mice; p<0.05*, p<0.01**, p<0.001***, p<0.0001**** two-tailed Student's t-test.

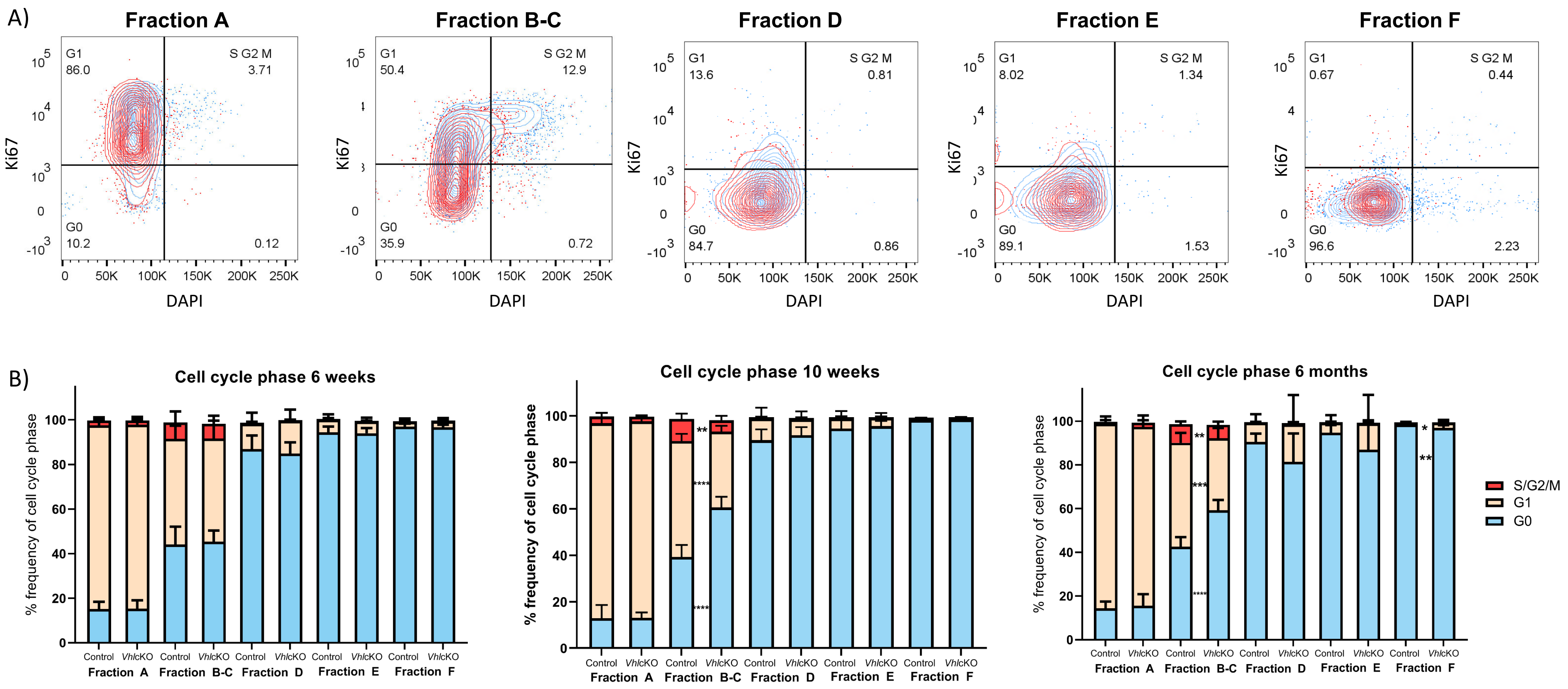


**Supplemental Figure 6. Proliferation analysis of B cell Fractions A-F** A) Representative FACS plot of cell cycle analysis in Fractions A-F cells (red: *Vhl*cKO blue: control); B) frequency of cells in each cell cycle phase within Fractions A-F at 6-weeks-old, 10-weeks-old old and 6-month-old mice. $p<0.05$*, $p<0.01$**, $p<0.001$***, $p<0.0001$**** two-tailed Student's t-test.

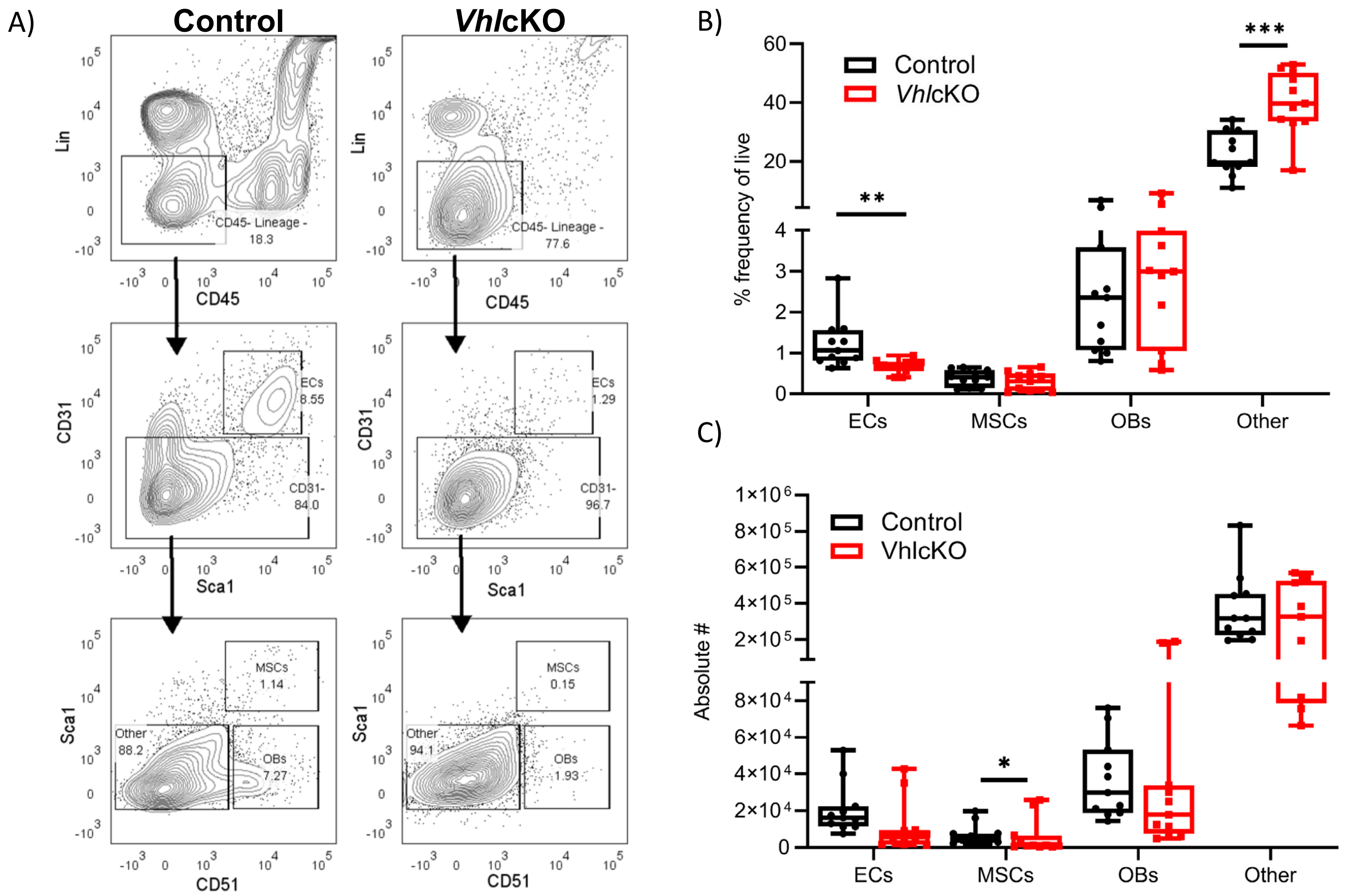


**Supplemental Figure 7. Flow cytometric analysis of *Vhl*cKO bones**. A) Flow cytometry gating strategy for bone niche cells (ECs, MSCs, OBs) after 2hr bone digest; B) frequencies and C) absolute numbers of each cell subset out of the live cell gate (not shown).

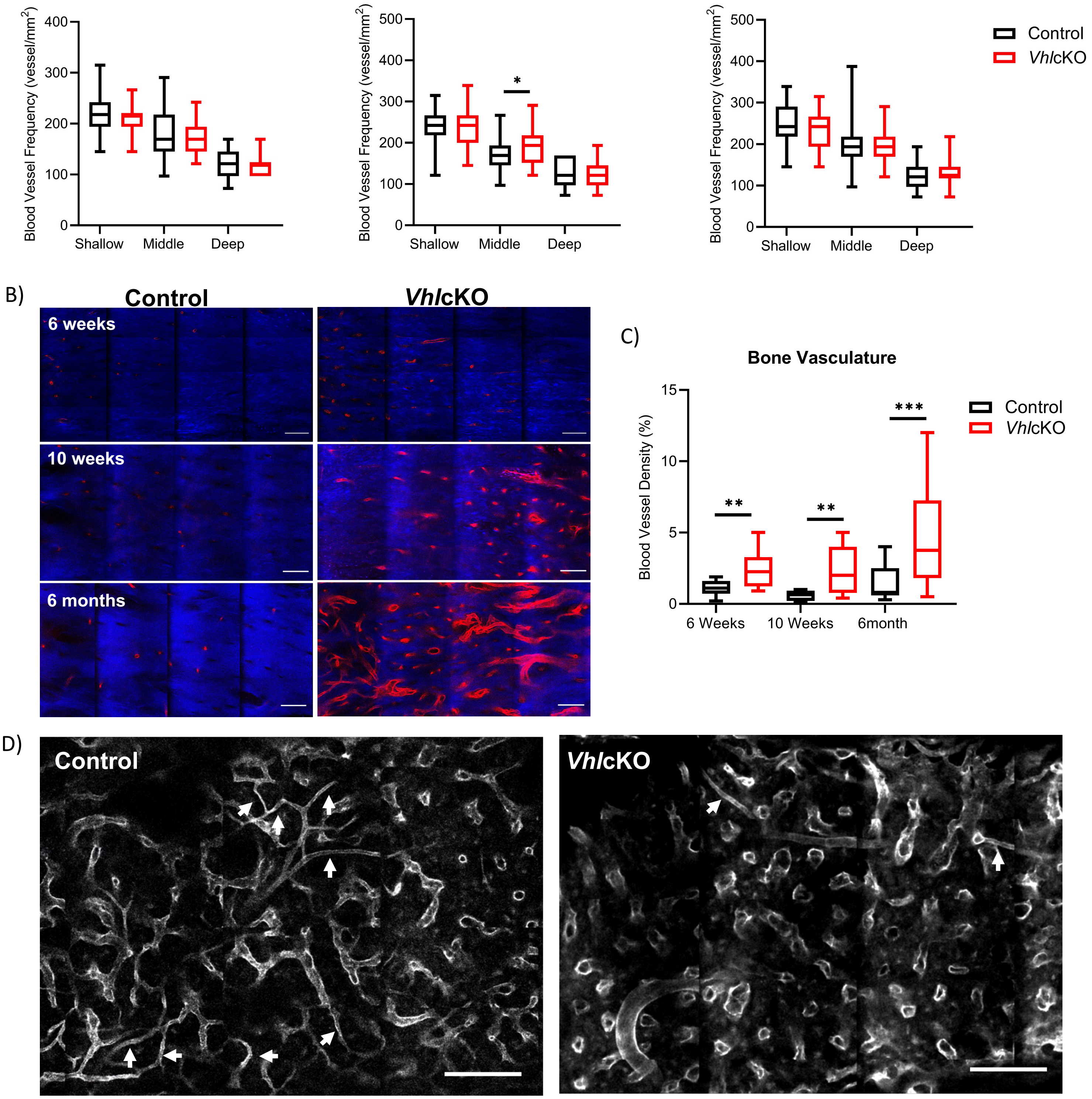


**Supplemental Figure 8**. **Ex vivo imaging of bone and bone marrow vasculature in *Vhl*cKO and control mice.** A) Quantification of age dependent vessel frequency in the diaphyseal BM of uDISCO cleared femurs at different depths (0-30 µm (shallow BM), 75-105 µm (middle BM), and 150-180 µm (deep BM) below the endosteum; B) maximum intensity projection images of cortical bone vascularization and C) quantification of cortical vessel density. Red: blood vessels (labeled with Alexa647 conjugated antibodies against CD31, CD144, and Sca-1), Blue: bone (SHG); D) maximum intensity projections of BM blood vessels within 50 µm of the endosteum in femurs of control (left) and *Vhl*cKO (right) mice; Grayscale: blood vessels (labeled with Alexa647 conjugated antibodies against CD31, CD144, and Sca-1). White arrows point to small diameter arterioles. Scale bars $\sim 100\ \mu m$.

**Supplemental Table 1.** List of the fluorochrome-labeled monoclonal antibodies used for flow cytometry and vessel staining (Alexa647)

| Antigen | Clone | Fluorochrome | Source |
|---|---|---|---|
| 7AAD | - | 7AAD | Biolegend |
| Annexin V | - | FITC | Biolegend |
| B220 | RA3-6B2 | PE-Cy7 | Biolegend |
| B220 | RA3-6B2 | APC-Cy7 | Biolegend |
| CD117 | 2B8 | APC-Cy7 | Biolegend |
| CD117 | 2B8 | APC-Cy7 | Biolegend |
| CD11b | M1/70 | APC-Cy7 | Biolegend |
| CD11b | M1/70 | biotin | Biolegend |
| CD11b | M1/70 | PE-Cy7 | Biolegend |
| CD127 | IL-7Ra | APC | Biolegend |
| CD135 | A2F10 | PE | eBioscience |
| CD150 | mShad150 | FITC | eBioscience |
| CD19 | 6D5 | PE | Biolegend |
| CD19 | 6D5 | FITC | Biolegend |
| CD19 | 6D5 | biotin | Biolegend |
| CD19 | 6D5 | PE-Cy7 | Biolegend |
| CD3 | 145-2C11 | APC | Biolegend |
| CD3 | 145-2C11 | biotin | Biolegend |
| CD3 | 145-2C11 | PE-Cy7 | Biolegend |
| CD31 | 390 | APC | Biolegend |
| CD4 | Gk1.5 | biotin | Biolegend |
| CD4 | Gk1.5 | PE-Cy7 | Biolegend |
| CD43 | 1B11 | PE | Biolegend |
| CD45 | 30F11 | FITC | Biolegend |
| CD45 | 30-F11 | PerCP Cy5.5 | Biolegend |
| CD45 | 30-F11 | BUV395 | BD Biosciences |
| CD45 | 30F11 | BV421 | Biolegend |
| CD45.1 | A20 | FITC | Biolegend |
| CD45.1 | A20 | BUV395 | BD Horizon |
| CD45.2 | 104 | APC-Cy7 | Biolegend |
| CD45.2 | 104 | APC | Biolegend |
| CD48 | HM48-1 | PE-Cy7 | Biolegend |
| CD51 | RMV-7 | Biotin | Biolegend |
| CD8 | 53.6.7 | biotin | Biolegend |
| CD8 | 53.6.7 | PE-Cy7 | Biolegend |
| Gr1 | RB6-8C5 | PE-Cy7 | Biolegend |
| Gr1 | RB6-8C5 | biotin | Biolegend |
| Gr1 | RB6-8C5 | PE-Cy7 | Biolegend |
| IgD | 12-26c.2a | BV510 | Biolegend |
| IgD | 11-26c.2a | FITC | Biolegend |
| IgG1,k | RTK2071 | Biotin | Biolegend |
| IgG2a,k | RTK2758 | APC | Biolegend |
| IgM | RMM-1 | BV421 | Biolegend |
| IgM | RMM-1 | PE-Cy7 | Biolegend |
| Ki67 | 16A8 | APC | Biolegend |
| Ly-6A/E | D7 | BV510 | Biolegend |
| Nk1.1 | PK136 | biotin | Biolegend |
| Nk1.1 | PK136 | PE-Cy7 | Biolegend |
| Streptavidin | - | Pacific Blue | Life Technologies |
| Streptavidin | - | PE | eBioscience |
| Ter119 | TER119 | biotin | Biolegend |
| Ter119 | TER119 | PE-Cy7 | Biolegend |
| CD144 | BV13 | Alexa647 | Biolegend |
| CD31 | MEC13.3 | Alexa647 | Biolegend |
| Ly-6A/E | D7 | Alexa647 | Biolegend |

#### SUPPLEMENTAL VIDEOS AND LEGENDS

**Videos 1-2. Representative ex vivo Videos recorded in the uDISCO cleared long bone BM**

Video 1. Representative 10-week-old Control uDISCO cleared long bone Z stack (scale bar ~200 µm)

Video 2. Representative 10-week-old Control uDISCO cleared long bone 3D view (scale bar ~100 µm)

**Videos 3-8. Representative Leakage Videos recorded in the calvaria BM**.

Representative zstacks of the calvaria BM recorded 10 minutes after Rhodamine B Dextran injection. Z step size is 2 µm and scale bars ~50 µm. Green Channel = bone (SHG), Blue = Rhodamine-B-Dextran (70 kDa). Brightness/Contrast adjusted for display only.

Video 3. Representative 6-week-old Control Leakage Zstack

Video 4. Representative 6-week-old *Vhlc*KO Leakage Zstack

Video 5. Representative 10-week-old Control Leakage Zstack

Video 6. Representative 10-week-old *Vhlc*KO Leakage Zstack

Video 7. Representative 6-month-old Control Leakage Zstack

Video 8. Representative 6-month-old *Vhlc*KO Leakage Zstack

**Videos 9-10. Representative permeability videos recorded in the calvaria BM.** Representative video of the calvaria BM permeability recorded immediately after Rhodamine-B-Dextran injection. Scale bars ~50 µm. Brightness/Contrast adjusted for display only.

Video 9. Representative 6-week-old Control Permeability Video

Video 10. Representative 6-week-old *Vhl*cKO Permeability Video
